# Deep learning reduces data requirements and allows real-time measurements in Imaging Fluorescence Correlation Spectroscopy

**DOI:** 10.1101/2023.08.07.552352

**Authors:** Wai Hoh Tang, Shao Ren Sim, Daniel Ying Kia Aik, Ashwin Venkata Subba Nelanuthala, Thamarailingam Athilingam, Adrian Röllin, Thorsten Wohland

## Abstract

Imaging Fluorescence Correlation Spectroscopy (Imaging FCS) is a powerful tool to extract information on molecular mobilities, actions and interactions in live cells, tissues and organisms. Nevertheless, several limitations restrict its applicability. First, FCS is data hungry, requiring 50,000 frames at 1 ms time resolution to obtain accurate parameter estimates. Second, the data size makes evaluation slow. Thirdly, as FCS evaluation is model-dependent, data evaluation is significantly slowed unless analytic models are available. Here we introduce two convolutional neural networks (CNNs) – *FCSNet* and *Im-FCSNet* – for correlation and intensity trace analysis, respectively. *FCSNet* robustly predicts parameters in 2D and 3D live samples. *ImFCSNet* reduces the amount of data required for accurate parameter retrieval by at least one order of magnitude and makes correct estimates even in moderately defocused samples. Both CNNs are trained on simulated data, are model-agnostic, and allow autonomous, real-time evaluation of Imaging FCS measurements.

---

The description of a biological system requires spatial and temporal information. While spatial information has reached its physical limits with Ångström resolution possible ^1,2^, temporal information on whole images is still not easily available. One method that can provide temporal resolution down to at least 40 µs with commercially available instrumentation is Imaging Fluorescence Correlation Spectroscopy, or Imaging FCS ^3,4^. Imaging FCS analyzes the fluorescence fluctuations in every pixel in a temporal image stack to determine the molecular processes underlying these fluctuations in the different parts of the sample. It can be implemented with either Total Internal Reflection Fluorescence (TIRF) microscopy, Variable Angle Illumination or Single Plane Illumination Microscopy (SPIM), either in vitro, in cells, or in vivo to provide parameter maps, such as concentrations, diffusion coefficients, or measures of interactions ^5^. Nevertheless, Imaging FCS has several limitations. Firstly, correlation functions are biased estimators, and a minimum of frames has to be taken to obtain accurate and precise parameter estimates ^6,7^. For typical characteristic process times of 10s of milliseconds, as commonly seen in Imaging FCS on cellular processes, about 1 minute total recording time is required ^8^. Secondly, the long measurements correspond to a large amount of data on the order of gigabytes, slowing data evaluation. Lastly, for some situations no analytical fitting models are available, such as SPIM-FCS (in preparation). In these cases, one needs numerical evaluations to fit the data, adding to the computational cost and making real-time evaluation difficult.

We propose deep-learning inspired Convolutional Neural Networks (CNNs) to overcome these challenges. CNNs are used widely in computer vision. Since the introduction of *AlexNet* ^9^ for image recognition, the growth of CNN applications has been exponential and has gone well beyond computer vision. CNNs are now used in various different domains, including speech recognition ^10^, language translation ^11^, and natural language processing ^12^. We have seen similarly inspired techniques adopted into scientific fields, and recent examples include physics ^13^, mathematics ^14^, and chemistry ^15^. In biology specifically, deep learning has been used to augment currently available solutions in a diverse range of applications, including medical image segmentation ^16^, drug discovery ^17^, super-resolution microscopy ^18^, single-particle tracking ^19^ and protein structure prediction ^20^. Deep feed-forward networks coupled with wavelet spectral analysis for noise detection were used in FCS two-color experiments ^21^. And machine learning methods, including CNNs, were applied to analyze FCS parameters for binary classification of cancer patients in their oncology research ^22^. A recent review ^23^ points towards the use of CNNs in addressing the challenges of FCS analysis.

Here, we build two CNN models, which we named *FCSNet* and *ImFCSNet*, that analyze the autocorrelation functions or the raw intensity traces, respectively, to estimate diffusion coefficients at all pixels of an image stack of a sample. We illustrate the construction of these models and their workflow of the data evaluation compared to nonlinear least-squares (NLS) fitting in Fig. 1. We use simulated data for training both CNNs, thus eliminating the cumbersome and difficult task of obtaining experimental training data for the required wide data range for the diffusion coefficient (D) and signal-to-noise ratio (SNR). Interestingly, we show that we can use the same architectures for 2D and 3D systems with different illumination modalities, i.e. TIRF or SPIM. We demonstrate the strength and weaknesses of *FCSNet*, and *ImFCSNet* by applying them to supported lipid bilayers, cells and drosophila melanogaster embryos. We show that depending on which CNN is used, we can estimate D from up to 20 times less data than what is required in traditional FCS measurements, parameter estimation can be made even when the sample is de-focused, and data evaluation can be sped-up by up to four orders of magnitude. In addition, both CNNs are fit model-agnostic and thus do not require functional forms for the autocorrelation function to extract D from a measurement, simplifying FCS analysis and broadening FCS applications to any desired illumination and detection geometry. Our new approach makes Imaging FCS a versatile and easy-to-use tool by providing diffusion coefficient maps across biological samples in real-time.

**Fig. 1.**
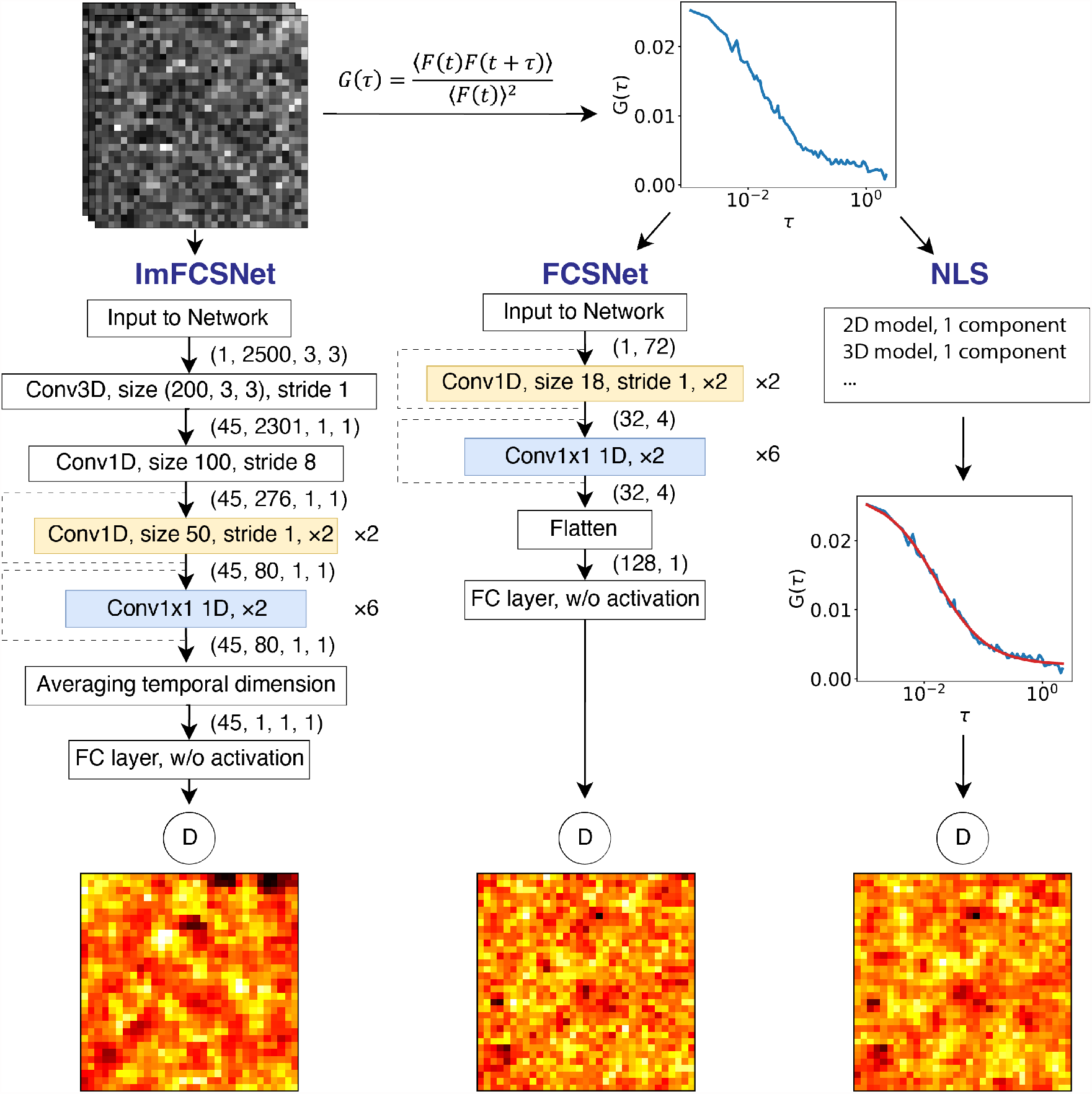
Schematic illustration of the CNN networks and the evaluation workflow. We base the CNN models on the ResNet architecture, where the skip connections are the dotted lines in the figure. The × 2 in the yellow and blue convolutional layers means these convolutional layers come in a pair within each residual block. We repeated the residual blocks two and six times, respectively. *ImFCSNet* evaluates 2,500 frames from the raw image stack through *ImFCSNet*. We can apply *ImFCSNet* in an overlapping manner to get a D map of individual pixels, excluding image edges. The ACF curves become the inputs for *FCSNet* and NLS curve-fitting evaluations.

## Results

We assess the CNNs on a variety of 2D and 3D samples, including supported lipid bilayers, cells, fluorescent bead solutions, and drosophila embryos (Supplementary Section 4), and a variety of different setup characteristics (Supplementary Table 1.1), to demonstrate the feasibility of applying the models to experimental measurements of different conditions. For each new setting, we retrain a CNN to ensure correct predictions. Since we do not have the ground truth of the measurements, we use NLS fits of a single diffusive component of the ACFs calculated from 50,000 frames of the experimental data as basis for comparison. The image stack is collected using custom-built data acquisition software, which enables real-time adjustments of the correlation function, thereby aiding in microscope alignment^24^. We evaluate the NLS fit models using the Imaging FCS ImageJ plugin ^25^, using standard Imaging FCS fit models, which are described in the Methods section. We can run the NLS evaluations with a workstation central processing unit (CPU) or can accelerate the process by leveraging the parallel computations on a graphic processing unit (GPU). Before any evaluation, we apply a polynomial bleach correction of order four, unless stated otherwise. Furthermore, we apply Z-score standardization to the input data of the CNNs, i.e. subtracting the mean and dividing by the standard deviation of the input data.

**Table 1.**
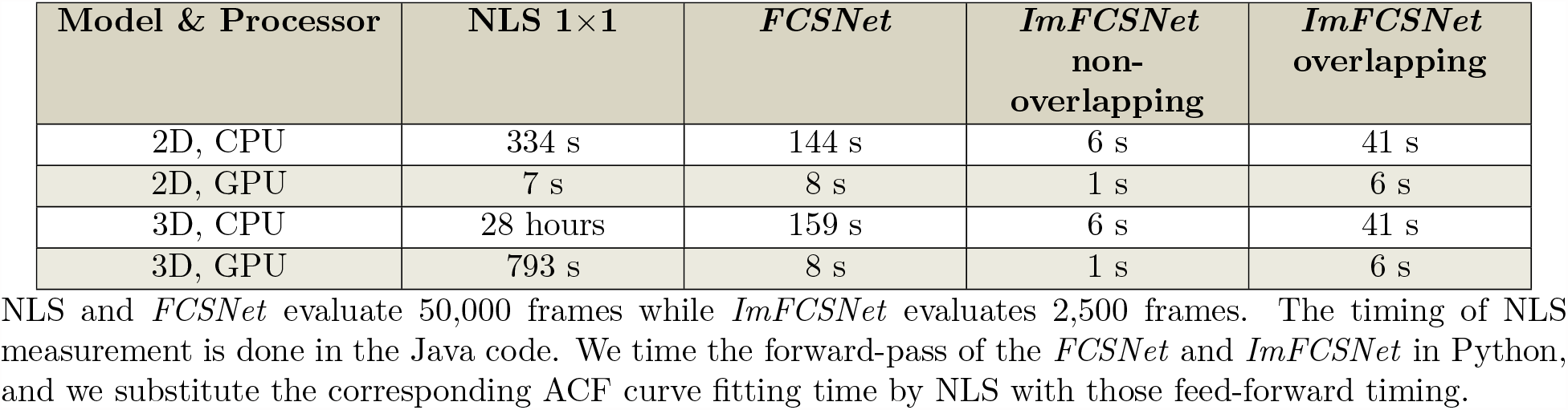
Evaluation speed.

| Model & Processor | NLS 1×1 | *FCSNet* | *ImFCSNet*<br>non-overlapping | *ImFCSNet*<br>overlapping |
| --- | --- | --- | --- | --- |
| 2D, CPU | 334 s | 144 s | 6 s | 41 s |
| 2D, GPU | 7 s | 8 s | 1 s | 6 s |
| 3D, CPU | 28 hours | 159 s | 6 s | 41 s |
| 3D, GPU | 793 s | 8 s | 1 s | 6 s |

NLS and *FCSNet* evaluate 50,000 frames while *ImFCSNet* evaluates 2,500 frames. The timing of NLS measurement is done in the Java code. We time the forward-pass of the *FCSNet* and *ImFCSNet* in Python, and we substitute the corresponding ACF curve fitting time by NLS with those feed-forward timing.

**Table 2.**
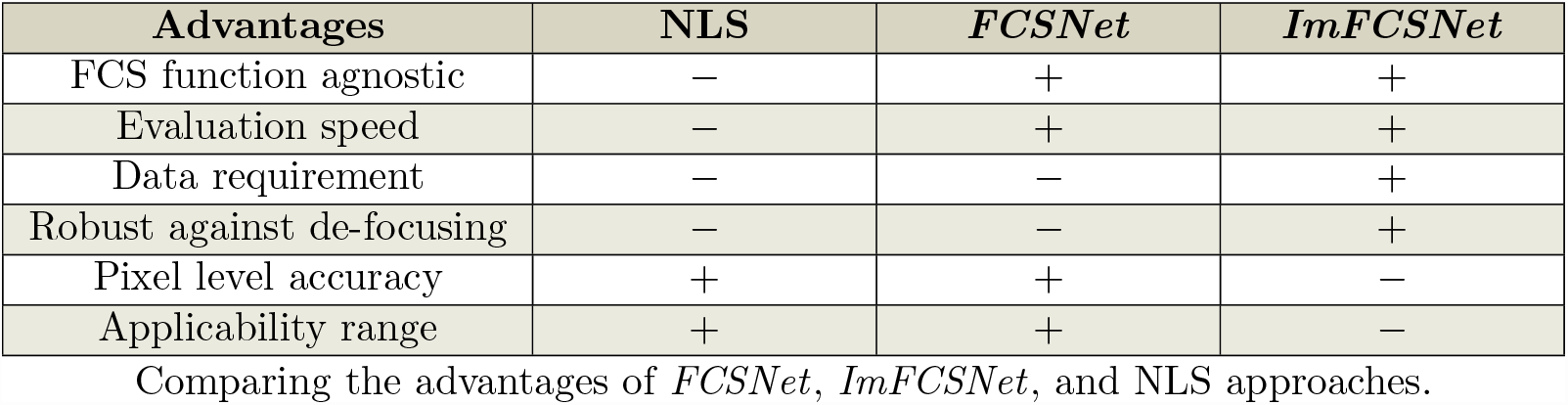
Summary of advantages.

A complete description of the two CNN models (Fig. 1) is in the Methods section and their training are in Supplementary Section 1. Briefly, the CNNs, based on the ResNet architecture ^26,27^, are trained on simulated data. For that purpose, we simulate random walks in 2D at constant illumination intensity (TIRF), or in 3D within a Gaussian Beam for a wide range of parameters at varying signal-to-noise ratio (SNR) settings (in Supplementary Tables 1.2 and 1.4). We assume a single diffusing particle in all cases. The CNNs therefore predict only one parameter, i.e. D. We target our model to evaluate a wide range of D, from 0.1 to 10 µm^2^/s, and we therefore let the model to output the natural log of D. *FCSNet* is trained on simulated ACFs calculated from 50,000 temporal data points as it will be applied to every single pixel of an image stack. We train *FCSNet* over 3,000 epochs and choose the model with the lowest training loss. *ImFCSNet*, on the other hand, is trained on 2,500 intensity values of groups of 3 × 3 pixels, generated on-the-fly. The training of *ImFCSNet* is essentially a 1-epoch training with many batches and the data is different for each batch. We have a training schedule (see Supplementary Table 1.6) where we train *ImFCSNet* for 8 rounds of the 1-epoch training, 48,000 batches per epoch. It is a progressive training where we increase the noise level of the training data in each round. Lastly, we also evaluate the trained *FCSNet* and *ImFCSNet* models on simulated test dataset (Supplementary Section 2).

### In vitro measurements

First, we tested a DOPC supported lipid bilayer labeled with Rhodamine–PE (0.01 mol% dye to lipid ratio). We collected 300,000 frames at a frame rate of 1.06 ms, which corresponds to 318 s or about 5 minutes of measurement time. The measurement was then partitioned into non-overlapping sets of either 2,500 (*ImFCSNet*) or 50,000 (*FCSNet* and NLS) frames for data evaluation. The network predictions for both CNNs were close to the values determined by NLS fits (Fig. 2a). While *FCSNet* achieves a better precision compared to *ImFCSNet*, it provides predictions only every 50 s, compared to every 2.5 s for *ImFCSNet* (Fig. 2b). Importantly, all three evaluations follow the same trend (Fig. 2b). The Spearman correlation coefficients between NLS and *FCSNet* and *ImFCSNet* are 0.89 and 0.60, respectively.

**Fig. 2.**
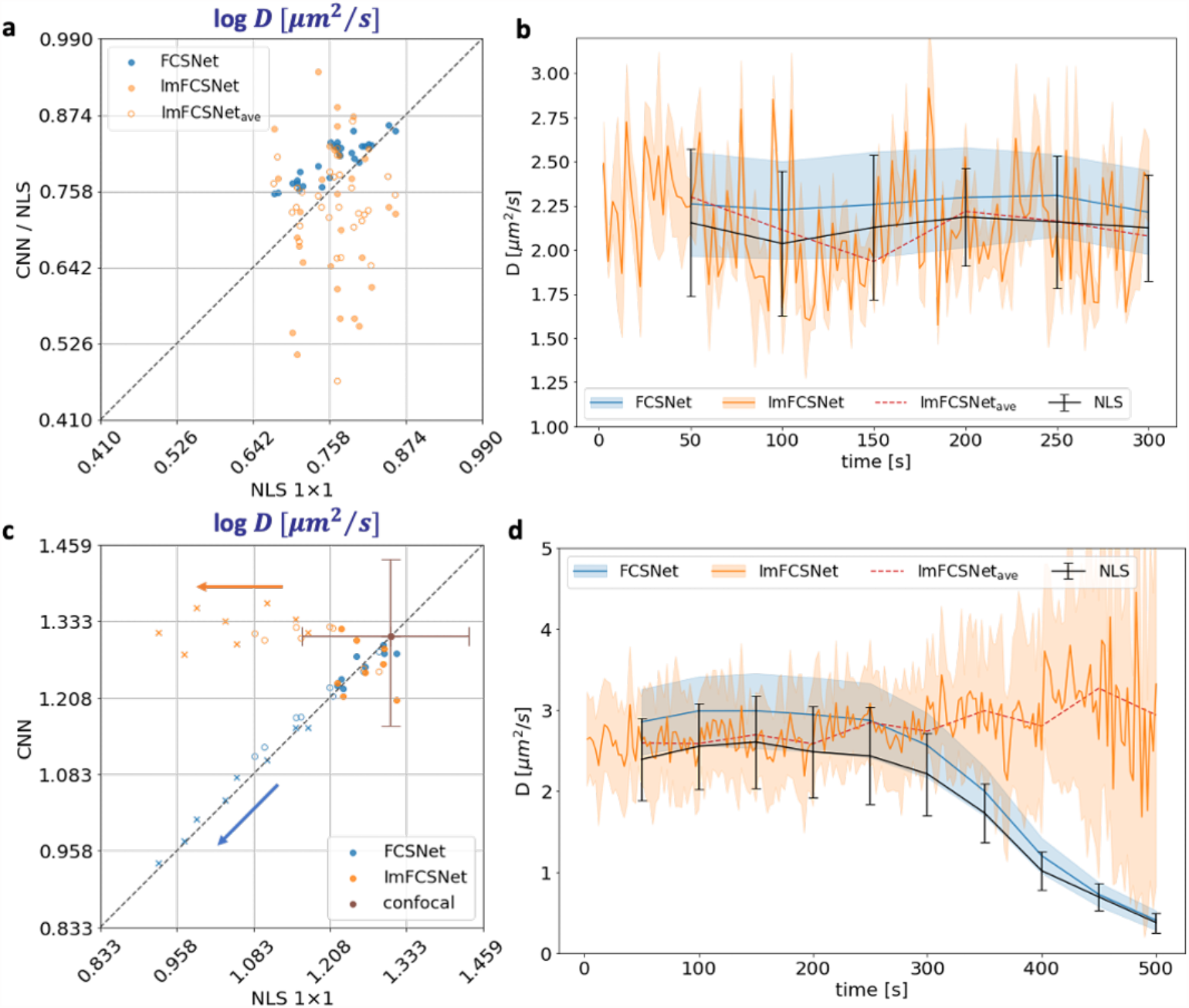
Evaluations on chemical samples. **a**, The average D values of multiple 6 × 6, 50,000 frames, ROIs in DOPC lipid bilayer measurements. The closer the points to the diagonal, the closer the predictions by the networks are to the NLS predictions. Averaging, i.e. *ImFCSNet*_*ave*_ evaluations, improves the predictions, as shown by the orange crosses. The range of standard deviations of the average diffusion coefficients for NLS, *FCSNet*, and *ImFCSNet* methods are 0.24 – 0.56 µm^2^/s, 0.20 – 0.39 µm^2^/s, and 0.09 – 0.75 µm^2^/s, respectively. **b**, The predictions of a 6 × 6 ROI region over 300,000 frames. *ImFCSNet* the dynamics within a shorter duration, which is shown by the evaluations done every 2,500 frames. The predictions typically show larger fluctuations but follow the same trend as NLS and *FCSNet* predictions. **c**, The average diffusion coefficients on bead measurements for in-focus and defocused measurements. The predictions, particularly the solid points, i.e. in-focus measurement, are distributed around the diagonal; thus, we get comparable networks’ to NLS predictions. The standard deviations of the average diffusion coefficients for NLS, *FCSNet*, and *ImFCSNet* methods are 0.46 – 0.67 µm^2^/s, 0.31 – 0.36 µm^2^/s, and 0.63 – 0.93 µm^2^/s, respectively. The confocal measurement, D of 3.7 ± 0.5 µm^2^/s, is an additional reference. *FCSNet* follows the same trend as NLS predictions when the measurements defocus. On the other hand, *ImFCSNet* predictions are more robust against defocused measurements. **d**, Evaluations of a 500,000 frames POPC lipid bilayer measurement become defocused over time. *FCSNet* follows the same trend as NLS when the measurement is defocusing, i.e. a dip in the diffusion coefficients. *ImFCSNet* predictions are relatively stable as the sample defocuses, for example, from 250 to 350 seconds. This characteristic is consistent with the results in **c**.

In the case of 3D measurements using SPIM-FCS, we used a sample of 100 nm fluorescent polystyrene beads in aqueous solution. As beads were highly fluorescent and did not show any decrease in intensity during the measurement, no bleach correction was applied. The CNNs’ predictions are consistent with the NLS fit. Moreover, the predictions fall within one standard deviation of a confocal FCS measurement, which serves as an additional reference point (Fig. 2c). Interestingly, when the sample is de-focused, the two CNNs show different behaviour. NLS fit and *FCSNet* evaluate the same ACFs and show the same trend, namely a decrease of D with increased de-focusing. Note that the values for *FCSNet* in Fig. 2c lie almost exactly on the diagonal, demonstrating how similar *FCSNet* predictions are to NLS fits. However, *ImFCSNet* predictions are robust against the change in focus as the values stay constant and are well within one standard deviation of the confocal measurement.

We conducted a similar test to determine whether *ImFCSNet* is also robust against de-focusing in the TIRF setup. Fig. 2d shows a TIRF-based Imaging FCS measurement of a POPC supported lipid bilayer over 500,000 frames (∼ 8 minutes), which was measured at a frame rate of 1.06 ms. It slowly de-focused from about the 250,000^th^ frame onward. *ImFCSNet* provides many more points and thus a better time resolution, and maintains consistent predicted D values throughout the measurement, unlike *FCSNet* and NLS. However, the standard deviation of the D values, predicted by *ImFCSNet*, increases as the SNR decreases with de-focusing.

### 2D measurements on live cells

CHO-K1 cells were transiently transfected with a plasma membrane targeted fluorescent protein, PMT-mEGFP^28^, and 50,000 frames at 2.06 ms frame rate of a 21 × 21 pixel area were recorded in TIRF. For a better and fairer comparison between NLS and the CNNs, we calculated the NLS fits for both, 1 × 1 and overlapping 3 × 3 binning and provide the D maps as well as the scatter plots of the CNNs against NLS (Fig. 3c). *FCSNet* produces a diffusion map that is similar to NLS 1 × 1, which is also reflected in the scatter plot Fig. 3c, where the points are mostly along the diagonals, and the standard deviation of NLS (with 1 × 1 binning) and *FCSNet* predictions are 0.39 and 0.47, respectively. *ImFCSNet* predictions, based on only 2,500 frames, are more widely distributed, in the scatter plot (Fig. 3f), as seen before also on the bilayer measurements. The standard deviation of NLS (with overlapping 3 × 3 binning) predictions in Fig. 3f is 0.32 while it is 0.73 for *ImFCSNet*. If we average the 20 predictions of *ImFCSNet*, denoted as *ImFCSNet*_*ave*_, we see a slight improvement in the precision of the evaluations, as shown by the reduction of the standard deviation of the predictions on the y-axis (Fig. 3i) to 0.69. It is nonetheless not as compact as the predictions of *FCSNet* (Fig. 3c).

**Fig. 3.**
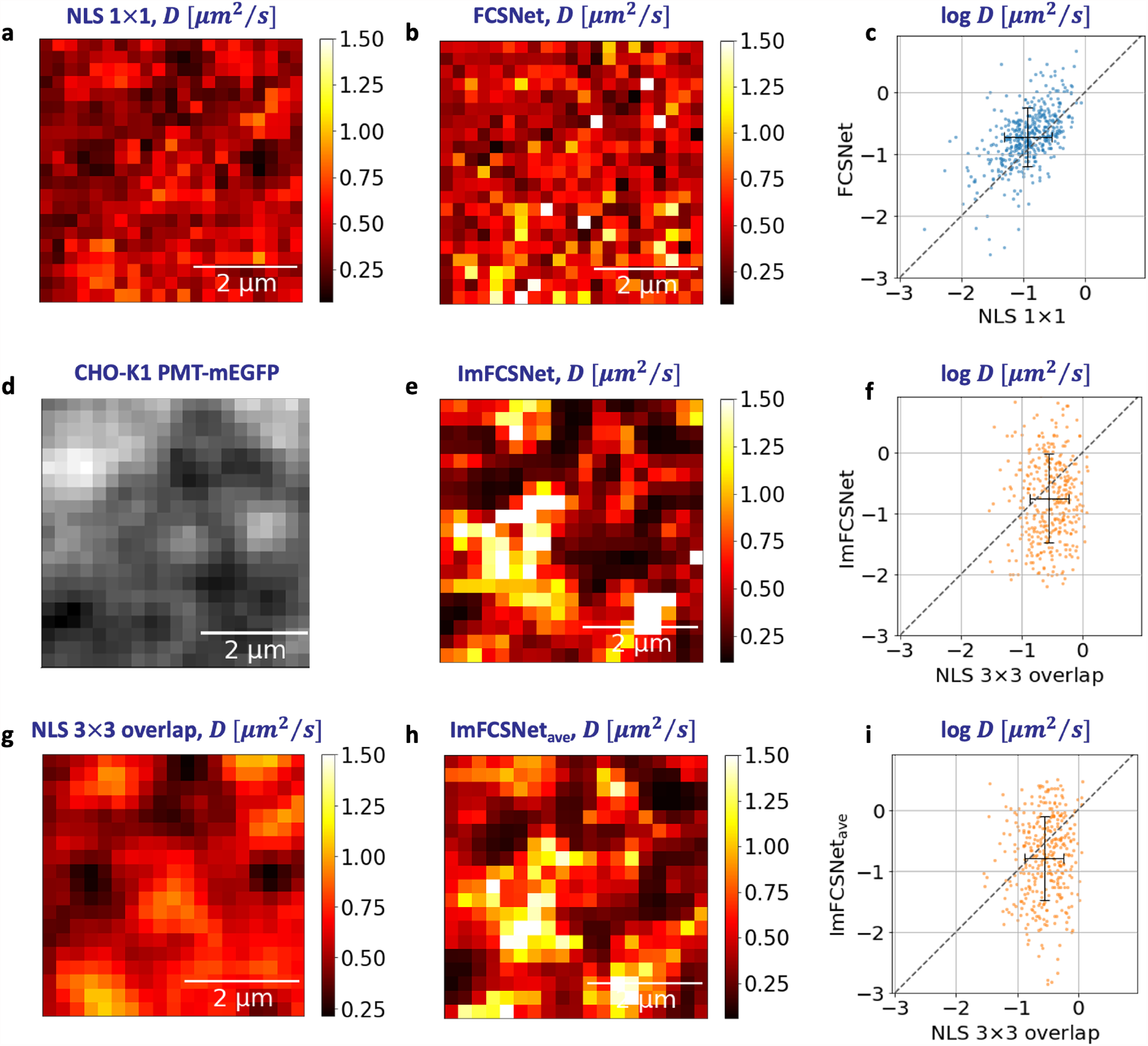
Evaluations on a CHO-K1 cell expressing PMT-mEGFP. **a**, NLS D map of 50,000 frames of individual pixels. **b**, *FCSNet* D map. **c**, Scatter plot of **b** vs. **a** predictions. **d**, An ROI of 21 × 21 pixels of a CHO cell expressing PMT-mEGFP. The widefield image is in Supplementary Fig. 6.1. **e**, *ImFCSNet* evaluation of the first 2,500 frames. **f**, Scatter plot of **e** vs **g** predictions. **g**, NLS D map with overlapping 3 × 3 binning of 50,000 frames. **h**, *ImFCSNet*_*ave*_: average of twenty *ImFCSNet* non-overlapping evaluations over the 50,000 frames. **i**, Scatter plot of **h** against **g** predictions. The black crosses in the scatter plots are the average and *±*1 standard deviation of the respective log D values.

### 3D measurements in an early drosophila embryo

Next, we used a transgenic eGFP-tagged bicoid (eGFP-Bcd) drosophila line ^29^. Bicoid is a morphogen that provides positional information to the syncytial nuclei of the embryo through their concentration gradient that forms across embryo’s anterior-posterior axis ^29^, a morphogen that regulates development in a concentration dependent manner ^30^, in drosophila embryos. Bicoid is known to be found inside and outside the nucleus where it can have various interactions ^31–33^. Here we recorded an image stack of 128 × 128 pixels and 50,000 frames at 2.04 ms per frame at the anterior domain (∼ 100µm from the anterior end) of the embryo at nuclear cycle (n.c.) 14. There is a slight drift observed in the measurement – over 4 pixels upward and 2 pixels to the right over 50,000 frames, for which we perform a simple drift correction by shifting 1 pixel downward every 12,500 frames and 1 pixel leftward every 25,000 frames. We cropped the image stack to a smaller 120 × 120 region for evaluation to avoid any border effects due to the drift correction. This is a much more challenging sample as it has lower SNR compared to the other measurements, shows sample drift, and positive ACFs cannot be seen at all locations. In addition, bicoid, due to its interactions, might not exhibit a pure one-component diffusion process ^34,35^. Therefore, it is a good test of the sensitivity of the CNNs to the input data. In Fig. 4 we indicate the locations at which NLS cannot fit the data as blue pixels and exclude them from further evaluations. Figs. 4a,b show the similarity in the outlines and values between the diffusion maps of *FCSNet* and NLS evaluations of individual pixels. The scatter plot in Fig. 4c confirms the observation and shows the correlation between NLS and *FCSNet* values. The diffusion coefficients measured are consistent with previous data of 0.3 µm^2^/s ^29^. More recent publications indicate that there are actually two different diffusion coefficients, one on the order of 1 µm^2^/s and one about 10 µm^2^/s ^34,36^. Our time resolution of 2.04 ms will not be sufficient to resolve the fast diffusion coefficient. Therefore, the NLS and *FCSNet* predictions are consistent with the slower diffusive component.

**Fig. 4.**
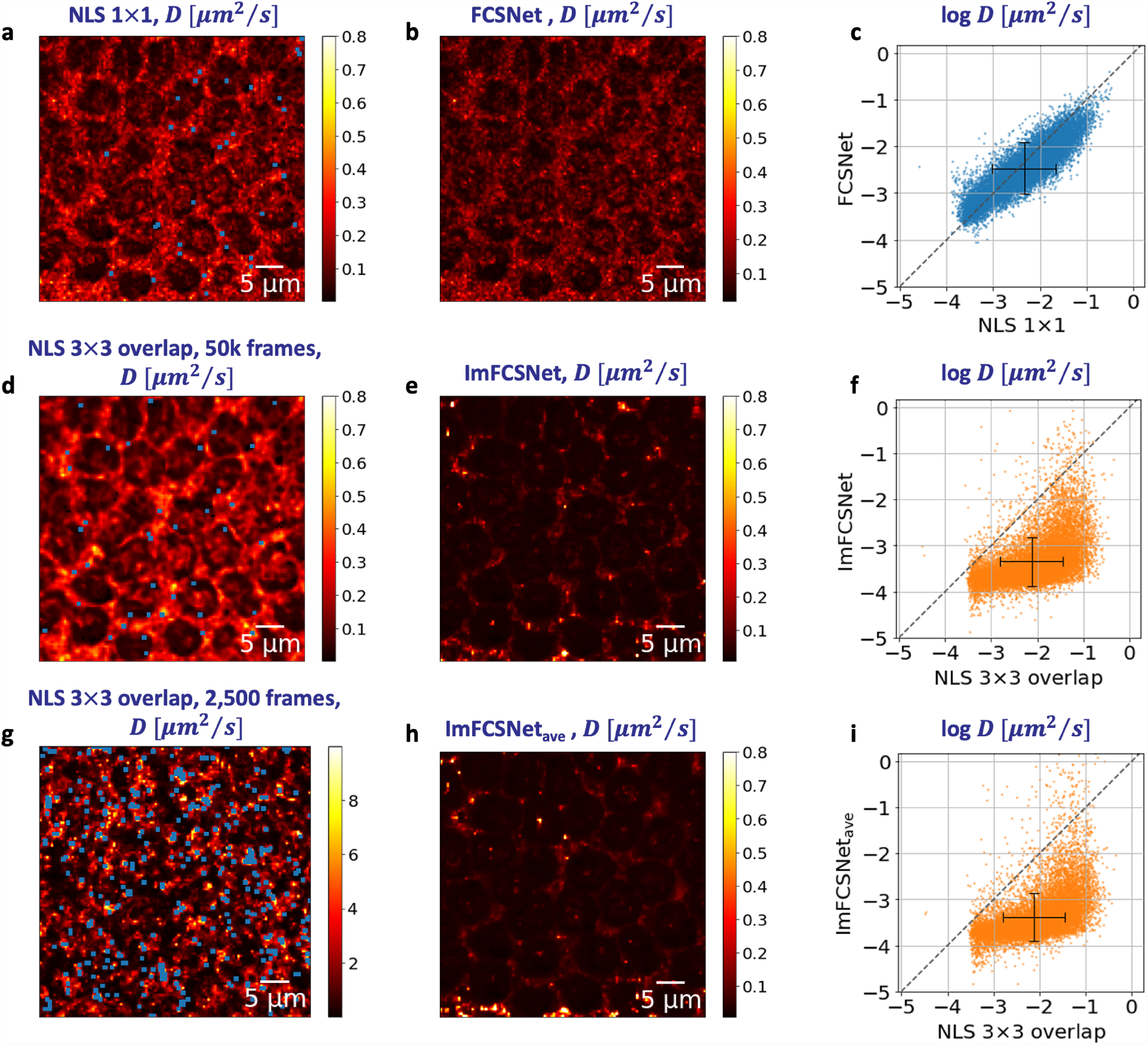
Drosophila embryo measurement on SPIM microscope. **a**, D map of NLS 1 × 1 evaluations for 50,000 frames, individual pixels. **b**, D map of *FCSNet* evaluations. **c**, Scatter plot of *FCSNet* vs NLS predictions. **d**, D map of NLS evluations for 50,000 frames with overlapping 3×3 binning. **e**, D map of *ImFCSNet* evaluations of the first 2,500 frames. **f**, Scatter plot of *ImFCSNet* vs NLS 3 × 3 with binning. **g**, D map of NLS evaluations for 2,500 frames, 3×3 overlapping binning. We observe that NLS at 2,500 frames predicts diffusion coefficients that are much higher than those for 50,000 frames in **d**. Even the genral sample outline cannot be discerned anymore. **h**, Averages of *ImFCSNet* predictions across time. **i**, Scatter plot of **h** against **d**. There is an increase in the diffusion coefficients for some pixels when we compute the averages. However, the NLS and *ImFCSNet* predictions still differ significantly. The tiny cyan squares in the diffusion maps are pixels that cannot be evaluated (or with large D values, which we capped at 10 µm^2^/s for this measurement) by the NLS, and they are excluded from our evaluations. The black crosses in the scatter plots are the average and *±*1 standard deviation of respective log D values.

For the comparison with *ImFCSNet*, we perform the NLS evaluation at overlapping 3 × 3 binning for 50,000 frames (Fig. 4d). In *ImFCSNet*, although the general outline of the sample is still visible in the diffusion coefficient map, the values are generally lower compared to NLS and *FCSNet* at 50,000 frames (Fig. 4e) and we observe skewed predictions in the scatter plot (Fig. 4f). Averaging multiple non-overlapping predictions of *ImFCSNet* across time did little to improve the results (Figs. 4h,i). However, it should be noted that *ImFCSNet* is a significant improvement over NLS at 2,500 frames (Fig. 4g), which does not preserve the sample structure in the diffusion coefficient map and can in general make no meaningful predictions, with diffusion coefficients overestimated by up to a factor 10.

These observations highlight the advantages and limitations of *ImFCSNet*. While *ImFCSNet* is a significant improvement over NLS at 2,500 frames, it does not reach the performance of *FCSNet*.

### Evaluation speed

Our networks offer a tremendous speed-up in evaluation time, offering (near) real-time analysis of measurements. The speed benchmarking is done on a workstation with an Intel(R) Core(TM) i9-10900KF CPU @ 3.70 GHz processor and a NVIDIA GeForce RTX 3090 GPU. We use the GPU implementation of the Imaging FCS plugin ^37^. With this newer generation of GPU, i.e. with more memory, cores and computing power, we observe in Table 1 a 47× speed-up for 2D NLS evaluations of 128 × 128 pixels, which is faster than the benchmark reported previously ^37^. For 3D NLS evaluation, the bottleneck in the processing time occurs as there is no analytic solution for the ACF fit function and numerical integration has to be used (equation (4)). While the GPU speeds up the NLS computation time by more than 100× compared to the CPU mode, it is still a time consuming process. Using CNNs, we improve the evaluation speed by another factor 100 for *FCSNet* and a factor 100 – 1,000 for *ImFCSNet* bringing evaluations to the second and thus real-time domain.

## Discussion

Imaging FCS is a powerful technique that provides information on biomolecular processes. Nevertheless, data evaluation can be complex and time-consuming, and several hurdles have hampered its widespread use. First, Imaging FCS measurements take much longer than simple imaging and require a recording of about one minute to obtain a sufficiently good signal-to-noise ratio. Although there have been recent attempts ^38,39^ to solve these issues, which could reduce measurement times to the range of seconds or even lower, they still have to show their applicability in vivo. Second, data fitting is strongly dependent on the availability of fit models. However, closed-form solutions for fit models are only available for simple measurement geometries and molecular processes. In other cases, numerical models can be used but require a much longer evaluation time. Third, the fitting process itself is complex and often requires expert users for interpretation ^40^. Fourth, FCS measurements strongly depend on the exact experimental conditions, and any misalignment will bias the results.

Here, we developed two CNNs to accelerate and simplify parameter estimation from single molecule fluorescence data. The first CNN, *FCSNet*, estimates the diffusion coefficient from ACFs of each pixel of a temporal image stack. The second CNN, *ImFCSNet*, reads in raw intensity traces directly from a 3 × 3 pixel area of a temporal image stack. We trained both CNNs on synthetic data. Here, we simulated exclusively free diffusion of one component. In principle, any process that changes fluorescence intensities and contributes to spatiotemporal ACFs can be simulated and used for training. The use of simulations for training solves two problems. First, it provides a wide range of ground truth data that can cover any desired range, while experimental data is limited by available standards with known diffusion coefficients. Second, the CNNs do not have to rely on theoretical ACF models as solutions for a particular process and thus become model-agnostic.

This has another important advantage. In this work, we introduced a numerical fit model for SPIM-FCS that takes into account the cross-talk from out of focus particles. The older SPIM-FCS models ^25^ underestimate D as they cannot account for the cross-talk. E.g. for the bead measurements in Fig. 2c, the older SPIM-FCS model predicts a D = 2.4 ± 0.4 µm^2^/s, which is 50% lower than the value of the new SPIM-FCS model at 3.5 ± 0.6 µm^2^/s. Our CNNs which are model-agnostic and were trained on simulations of the actual situation provide predictions of D much closer to our numerical model (*FCSNet* : 3.5 ± 0.4 µm^2^/s; *ImFCSNet* : 3.5 ± 0.8 µm^2^/s), thus justifying it. Finally, the value of the same sample in confocal FCS is 3.7 ± 0.5 µm^2^/s confirming the larger D value. Thus CNNs do not only provide parameter predictions but can validate theoretical models.

The CNNs can provide 1 – 4 orders of magnitude faster data treatment than NLS fitting, depending on the particular situation. The data passes through a single evaluation step in a CNN compared to NLS, which iteratively evaluates fits until it reaches a user-defined precision goal. As *FCSNet* requires the calculation of ACFs, similar to NLS, it reduces the data evaluation time to a lesser extent than *ImFCSNet*, which directly uses the raw intensity stacks as input. Therefore, *FCSNet* reduces data evaluation times between a factor 2 (CPU, 2D data with analytic fit model) and a factor 600 (CPU, 3D data with numerical fit model). On the tested GPU, this gain reduces and provides only improved performance for the 3D case with a numerical fit model, where the gain is about a factor of 100. *ImFCSNet* improves evaluation times for the same data by about 1 – 4 orders of magnitude on the CPU and about 1 – 3 orders of magnitude on the GPU. As can be seen, the speed-up is particularly advantageous when no analytic solutions for the ACF are available (Table 1). Furthermore, the CNN analysis is essentially real-time and does not require user input, thus making Imaging FCS available to non-expert users.

The two CNNs have their unique advantages. *FCSNet* achieves similar, if not better, accuracy and precision compared to NLS in 2D and 3D measurements. It evaluates the data of single pixels but requires, at the moment, still 50,000 frames. *ImFCSNet*, which evaluates the raw data and not the correlation functions, has additional advantages. It can obtain similar results as NLS with up to 20 times fewer data, making FCS measurements shorter and allowing to follow biomolecular processes not accessible up to now due to the long measurement times of FCS. Furthermore, *ImFCSNet* is less sensitive to focusing and provides stable parameters even when the sample is defocused. We speculate that the access to spatial information through the 3 × 3 binning allows *ImFCSNet* to correct for defocusing, an advantage not available to *FCSNet*, which works on data from single pixels. The advantage of spatial information in *ImFCSNet* might also be used to correct aberrations when measuring in biological tissues, functioning essentially as a software-based adaptive optics approach. However, this requires further investigation.

The advantages of *ImFCSNet*, though, come at the price of reduced spatial resolution as it evaluates 3 × 3 patches of pixels. And although *ImFCSNet* can be run in an overlap mode and thus provide diffusion coefficient estimates at each pixel, the estimates of neighbouring pixels will be strongly interdependent. Nevertheless, even when using the same 3 × 3 areas, NLS cannot reach the same performance due to the bias of the ACFs at short measurement times of about 2.5 s (i.e. 2,500 frames), which cannot be entirely overcome by pixel binning.

On the flip side, CNN models also have drawbacks. The largest is the lack of explainability of how the models reach a particular decision. It was one of the motivations for developing *FCSNet*, which uses less than 100 points as input as it directly evaluates correlation functions. We speculate that the smaller CNNs with fewer parameters reduce the complexity of understanding a CNN, leading to clearer design principles when applying a single CNN structure and translating it between different parameter ranges, a topic to be addressed in the future. In addition, *ImFCSNet* predictions are sensitive to the SNR. We included supporting evidence in Supplementary Section 8 showing the larger fluctuations in *ImFCSNet* predictions for measurements taken at a low laser power and, thus, lower SNR. Furthermore, we have seen in the drosophila measurement in Fig. 4, that *ImFCSNet* fails. While it is an improvement over NLS at 2,500 frames and can at least retain some of the diffusion map features of the sample, it cannot reach the performance of NLS and *FCSNet* at 50,000 frames. In this case the evaluation of the ACF by NLS and *FCSNet* over 50,000 frames provides a more robust readout and the short measurements of *ImFCSNet* of 2,500 frames at moderate SNR are not sufficient to retrieve the correct diffusion coefficients. In addition, recent publications indicate that for bicoid there is a second fast diffusive component (∼ 10 µm^2^/s) that Imaging FCS cannot capture due to the limited time resolution of 2.04 ms ^29^,36. Thus our FCS analysis provides only the slower diffusive component of ∼ 1 µm^2^/s. However, the fast diffusive component could still contribute to the dynamics seen over 3 × 3 pixels of *ImFCSNet*, something not included in the training set.

We can explore different approaches to tackle these shortcomings. From a modelling perspective, we can experiment with Bayesian approaches ^41,42^ to determine a network’s uncertainty when evaluating out-of-distribution samples. From a data perspective, future additions of multiple components into the CNN training and faster measurements could resolve the issue of multiple components. The failure of *ImFCSNet* on this sample is a reminder that one must carefully determine in what range of experimental settings a CNN can operate reliably. Unlike NLS, which is statistically well-understood and provides with the 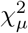 statistics a measure of the quality of fit, this is missing for CNNs. While CNNs can provide many advantages over NLS as discussed earlier, they can as well fail and that failure prediction is at the moment not possible. The drosophila experiment is a case in point where one CNN, *FCSNet*, works very well while the other, *ImFCSNet*, fails. One way to avoid overconfidence in a single prediction is to exploit the fact that our two CNNs have different advantages and drawbacks and could be used together. Due to their fast evaluation, once trained, the two or more models could be used in conjunction even for real-time evaluations. This would provide a mutual quality control that could be further supported by NLS checks. For instance, when using *FCSNet* and *ImFCSNet* simultaneously, they combine their advantages and are model-agnostic, provide significantly faster data evaluation compared to NLS fitting, are less sensitive to de-focusing artefacts or PSF changes, require fewer data to obtain the same parameters compared to standard Imaging FCS, and work in 2D and 3D on a wide range of samples, including supported lipid bilayers, cells, and multicellular organisms. We summarize the benefits of our models in Table 2. Spot checks by NLS would then ensure overall consistency of results.

While this is only a starting point, the advantages already shown by this first approach demonstrate the possibilities of CNNs to provide real-time data evaluation of experiments that are fast, model-agnostic, and robust against experimental problems, e.g. de-focusing. Future work will estimate more parameters, consider more heterogeneous systems with more than a single component and a wider range of processes, including more complex diffusion processes or active transport, and exploit the capability of CNNs to provide robust parameter estimates even in the case of non-ideal experimental conditions. Finally, the CNNs do not require input from expert users and thus make Imaging FCS readily available for a broad base of researchers.

## Methods

### Imaging FCS

At the heart of Imaging FCS is the fitting of measurements to a theoretical model based on the molecular properties of the fluorophores, the illumination profile, and the detection efficiency of the microscope setup. The workflow to obtain fitted parameters in the theoretical model involves several computations. Firstly, we compute the statistics of an input image stack in the form of the temporal autocorrelation function (ACF) and its standard deviation at each pixel of the image stack, where the ACF is given by

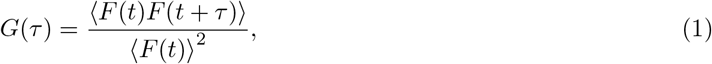

where *F(t)* is a measure of the fluorescence intensity at time t measured in a pixel and *τ* is the lag time between two time points in an image stack. The value of *F(t)* will be a function of the illumination intensity within the sample, the absorption cross-section and quantum yield of the fluorophore, the detection efficiency and the point spread function (PSF) of the microscope setup, and the distribution of fluorophores in the sample ^40^.

As the intensity traces in the image stack are temporally correlated, we use a blocking transformation to obtain unbiased estimates of the standard deviations ^43^. The reciprocals of these standard deviations act as weights in a non-linear least squares (NLS) fit ^44^ using the Levenberg-Marquardt algorithm ^45,46^. The ACF fit will provide the diffusion coefficient of the measured molecules at the location of a pixel, assuming a single molecular species in the sample, and the average number of molecules *N* within the observation area (2D) or volume (3D) of a pixel. For all our analysis, we compute the ACF based on correlators P of 16 and Q of 8. Correlator P is the number of correlation channels for the first group of bin times. The bin time is equal to the frame time / acquisition time. Correlator Q is the number of groups of bin times including the first). The bin time within a group is constant but it will always be twice as large as the bin time of the preceding group.

### 2D free diffusion

The ACF for free diffusion in 2D of a single molecular species ^47^ is

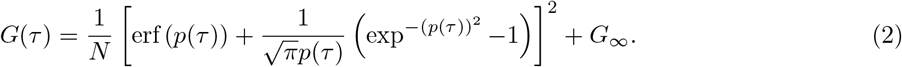

The term *p*(*τ*) is

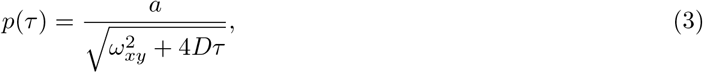

where *a* is the camera pixel pitch, *ω*_*xy*_ is the 1/*e*^2^ radius of a 2D Gaussian which is used to approximate the microscope PSF, *G*_*∞*_ is the convergence term of *G*(*τ*) as *τ* → ∞. The expected value of *G*_*∞*_ is 0 for long lag times and infinite measurement time. We use it, however, as a fit parameter as it can deviate slightly from 0 due to the finite measurement times. We recover the diffusion coefficient *D* by fitting an ACF curve to equation (2).

### 3D free diffusion

The ACF equation for Imaging FCS for 3D free diffusion of a single molecular species (in preparation) is given by

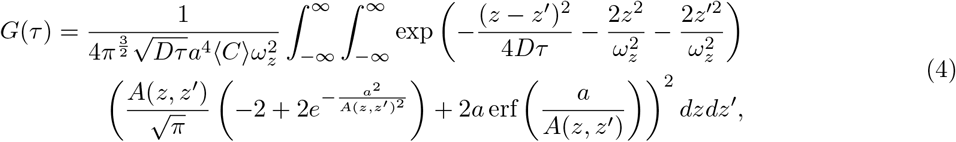

Where

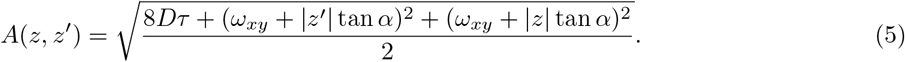

*a* is the maximal half-angle of the cone of light that enters/exits the objective lens, which can be computed given the numerical aperture of the lens and the index of refraction of the medium. *ω*_*z*_ is the 1/e^2^ radius of the light sheet thickness, and !_*xy*_ is the 1/e^2^ radius of the PSF of the objective in the x-y plane. There is no closed-form expression for equation (4). We require numerical evaluations for the double integral term, which adds to the computational cost when fitting the equation. We want to highlight the term *ω*_*xy*_ + z tan ff, which accounts for cross-talk between pixels from out-of-focus particles in the z-direction. It is an additional term to the 3D diffusion equation ^48^ and we incorporated it in the simulation of training data for our CNN models.

### Convolutional Neural Networks (CNNs)

Both *FCSNet* and *ImFCSNet* (Fig. 1) are inspired by the ResNet architectures ^26,27^, where the key feature is the skip connection that feeds the output of one layer as an input(s) to the next layer(s). Skip connections are also known as shortcut connections or identity mapping. They are depicted by dotted lines in Fig. 1. Skip connections have been shown to improve the stability of training, and there are different theories attempting to explain this observation, for instance, they smooth of the loss landscape ^49^, and they reduce the shattered gradients problem ^50^. The skip connections also allow the formation of ensembles of relatively shallow networks in a residual network ^51^. Putting it differently, the skip connections allow multiple connections between different blocks in the network and each block can be perceived as a smaller network. Our models, *FCSNet* and *ImFCSNet*, share many similarities in their construction. Every convolutional layer is followed by a batch normalization layer before applying ReLU activation. We have two residual blocks in the body of the networks, and each block is made up of two 1D convolutional layers and one skip connection. We do not use padding in our convolutional layers. Instead, we truncate the end of the skip connection to match the residual block’s temporal dimension and perform element-wise addition. We also deploy 1 × 1 convolutional layers ^52^, i.e. Conv1×1 1D layers. Similarly, we connect a skip connection to every two 1 × 1 convolutional layers. There are six such blocks in total. We included blocks of Conv1×1 1D layers to increase the depth of the network thus allowing the information to flow through more paths within the network.

There are three main differences in the two models. The first difference is at the input layer. The input to *FCSNet* is an ACF curve of 72 points, while the input to *ImFCSNet* is a raw image stack of 2500 frames of 3-by-3 pixels. We perform Z-score standardization to the input by subtracting its mean and dividing by its standard deviation. The input of *FCSNet* goes directly from the input layer to a 1D convolutional layer. On the contrary, we apply a 3D convolutional layer to the spatiotemporal input in *ImFCSNet*. It has a filter size of (200, 3, 3), stride of 1 and gives an output of 45 feature maps. This layer reduces the 3D input to 1D temporal data. Furthermore, we apply a 1D convolutional layer with a filter size of 100 and a stride of 8 in the second layer. We use a large stride in the second layer to reduce the memory footprint as data traverse through the network and to speed up the training process because there are fewer convolution computations in later layers.

We connect the output of the last Conv1×1 1D layer to a fully connected (FC) layer. The second difference between *FCSNet* and *ImFCSNet* here is how the output of the Conv1×1 1D is flattened. In *FCSNet*, we apply a standard flattening layer, which allows the data to match the dimension of the FC layer. In *ImFCSNet*, we perform an averaging along the temporal dimension of all feature maps. The averaging is equivalent to a global average pooling of each feature map ^53^. Some benefits of using an averaging operation are 1) the flexibility to pass in different length of input data and 2) experimenting with different Conv3D filter size and/or strides when training with more number of frames, without changing the number of layers in *ImFCSNet*. The last layer, i.e. the FC layer, does not apply an activation function and essentially performs a linear combination of the Conv1×1 1D output. Finally, the networks output the log of the diffusion coefficient. The last difference is the fewer feature maps in *FCSNet*. All in all, our models are lightweight: *FCSNet* and *ImFCSNet* contain 70,849 and 716,896 parameters, respectively. The forward pass is consequently fast. We included our training details in Supplementary Section 1.

## Data Availability

The data that support the findings of this study are available from the corresponding author, TW, upon reasonable request.

## Code Availability

The Python code has been publicly released under a MIT license and is available at https://github.com/waihoh/ImFCS_FCSNet_ImFCSNet.

## Acknowledgments

TW gratefully acknowledges funding by the Singapore Ministry of Education (MOE2016-T3-1-005 and C-33-18-279-V12). We are grateful to Timothy E. Saunders for discussions and the drosophila embryos.

## Author contributions

TW and AR conceived and oversaw the project. WHT and SRS created the CNNs. DA performed TIRF-FCS measurements, and adapted the ImFCS plugin for the current measurements. TA prepared the drosophila embryos. AVSN developed the new SPIM-FCS model and performed the drosophila embryo measurements. All authors wrote and agreed on the manuscript.

## Additional Information Supplementary information

The online version contains supplementary material.

