## supplementary material for "Deep learning reduces data requirements and allows real-time measurements in Imaging Fluorescence Correlation Spectroscopy"

August 7, 2023

#### 1 Training of CNNs

We used simulated training data of 2D or 3D free diffusion of fluorescent particles to train *FCSNet* and *ImFCSNet*. We add (camera) noise to the simulated data by sampling from either a Gaussian noise with varying variance or an empirical EMCCD noise distribution. The EMCCD noise data distribution is generated from a dark image measurement on the EMCCD camera, and Hirsch correction[54] was applied. We developed our models using the TensorFlow v2.5 deep learning library and trained them on NVIDIA RTX 3080 GPUs. We used Adam optimizer in our training. Once trained, we target our CNNs to evaluate 1-particle free diffusion in the range of 0.1 to 10 $\mu m^2/s$.

##### 1.1 Simulation of image stacks

Simulation of Imaging FCS experiments are based on the algorithm described elsewhere[7,55]. We make a minor change by converting the experimental units from *meters* and *seconds* to dimensionless units, *pixels* and *frames*. This renders the simulation parameters generic and easily adaptable to different setups. Given the initial experimental settings, we can easily find the corresponding predicted values in actual physical units. We first define the size of the pixel in sample space $\tilde{a}$ to help us convert the length unit to the *pixel* unit:

$$\tilde{a} = \frac{a}{\text{objective magnification}}, \tag{6}$$

where $a$ is the size of a camera pixel, for example, 24 $\mu m$. The objective magnification in our experiments are 60× for SPIM and 100× for TIRF. The parameters in the simulations are

$$\begin{aligned}
\sigma_{particle} &= \sqrt{2 \times D \times t_{step}} \times \frac{1}{\tilde{a}}, \\
\omega_{xy} &= \frac{\omega_{xy0} \times \lambda_{em}}{2 \times NA} \times \frac{1}{\tilde{a}}, \\
\rho &= \frac{N_{sim}}{A_{sim}}, \\
np_{step} &= CPS \times t_{step}, \\
\sigma_{ccd}^2 &= \text{variance of Gaussian noise}, \\
z_f &= \tan \arcsin \frac{NA}{n_{water}}, \\
\omega_z &= \frac{\omega_{z0} \times \lambda_{em}}{2 \times NA} \times \frac{1}{\tilde{a}},
\end{aligned} \tag{7}$$

where $D$ is the diffusion coefficient, $t_{step}$ is the time per simulation step, i.e. frame time $t$ divided by the steps per frame, $\omega_{xy}$ is the $1/e^2$ radius of a 2D Gaussian to approximate the microscope PSF (in x-y plane), with $\omega_{xy0}$ an experimentally determined constant. The parameter $\lambda_{em}$ is the emission wavelength, $NA$ is the numerical aperture of the microscope objective, and $CPS$ is the photon counts per particle per second. $\rho$ is the particle density, $N_{sim}$ is the the number of particles in the simulation area, $A_{sim}$, $np_{step}$ is the average number of photons emitted by a particle during the time $t_{step}$, $z_f$ is a factor used to calculate the spread of the PSF cross-section on the camera if the particle is not in the focal plane for 3D simulations. $n_{water}$ is the refractive index of water, which is 1.333. $\omega_z$ is the $1/e^2$ radius of the light sheet thickness, with $\omega_{z0}$ an experimentally determined constant. Supplementary Table 1.1 shows the different experimental settings for corresponding measurements.

**Supplementary Table 1.1 :** Different experimental settings

| **Parameters** | **DOPC & POPC lipid bilayers (2D – Setting 1)** | **CHO-K1 PMT-mEGFP & HeLa GPI-GFP (2D – Setting 2)** | **100 nm beads & drosophila embryo (3D)** |
|---|---|---|---|
| $a$ ($\mu m$) | 24 | 24 | 24 |
| objective magnification | 100× | 100× | 60× |
| $\lambda_{em}$ (nm) | 583 | 507 | 515 |
| $\omega_{xy0}$ | 0.8 | 1.01 | 1.1 |
| $\omega_{z0}$ | – | – | 2.2 |
| $NA$ | 1.45 | 1.49 | 1.0 |
| $t$ (ms) | 1.06 | 2.06 | 2.04 |

Summary table of different experimental settings.

We will now describe our simulation process, which is also summarized in Algorithm 1. Let our simulation area be $A_{sim}$. Our ROI is located at the center of $A_{sim}$. In our simulations, our ROI size is fixed at 3×3 pixels. There is a margin around the ROI. For example, if our simulation area is 15×15 pixels, then the margin is 6 pixels on each side of the ROI. For 2D diffusion, the particles will diffuse on the $x$-$y$ plane, while in 3D diffusion, the particles can diffuse in the $z$ dimension as well. The length of the simulation volume along the $z$ dimension is, 20× $\omega_z$ pixels, i.e. 10× $\omega_z$ pixels on each side of the focal plane. The $x$, $y$, and $z$ coordinates are in term of pixels. $N_{sim}$ is the number of diffusing particles. These $N_{sim}$ particles are positioned randomly in the simulation area for 2D simulation or in the simulation volume for 3D simulation at the beginning of the simulation. Each particle undergoes a Brownian motion in the free diffusion simulation. To determine the change in position of the $i$-th particle, we sample a random number from a standard normal distribution and multiply with $\sigma_{particle}$ to obtain the displacement along $x$ dimension at a time step $t$, i.e.

$$x_t^i = x_{t-1}^i + \sigma_{particle} \times N(0,1). \tag{8}$$

The subscript denotes the time step, superscript $i$ denotes the $i$-th particle, and $N(0,1)$ means a random sampling from a standard normal distribution. We compute in the same manner for $y_t^i$ dimension (and $z_t^i$ dimension in 3D simulation).

The number of photons, $N_{photon}$, emitted by each particle is sampled from a Poisson distribution with mean of $np_{step}$. Moreover in 3D simulation, the number of photons detected decays exponentially in $z$ direction, i.e. $\lfloor N_{photon} \times \exp(-\frac{1}{2}(\frac{z}{\omega_z})^2) \rfloor$. Given the ($x_t^i$, $y_t^i$) coordinates of the $i$-th particle, the position of each the photon is computed as

$$\begin{aligned} x_{photon} &= x_t^i + \omega_{xy} \times N(0,1), \\ y_{photon} &= y_t^i + \omega_{xy} \times N(0,1). \end{aligned} \tag{9}$$

We increase the photon count at each pixel in the ROI if the photon falls the ROI. For 3D simulation, we need to account for the increased size of the PSF cross-section on the camera, when the particle is not in the focal plane. Given the position $z$ of the particle, we approximate the PSF as

$$\tilde{\omega}_{xy} = \omega_{xy} + |z| \times z_f, \tag{10}$$

and replaces the PSF, i.e. the $\omega_{xy}$ term, in equation (9).

If the particle moves out of the simulation area or volume, we will place the particle at the boundary of the simulation area or volume randomly. This keeps the density of the particles in the simulation area unchanged. To take account of the movement of particles during the acquisition, we simulate multiple steps per frame and divide $t$ by the number of steps per frame. In our simulations, we choose 10 steps per frame. We accumulate the photons detected in all steps to create one frame. Finally, we can include camera noise to our simulation by sampling from a Gaussian distribution with variance $\sigma_{ccd}^2$ or from an empirical EMCCD noise distribution for each pixel in each frame. To generate a wide range of simulated data, we sample a range of simulation parameters, which are described in Supplementary Tables 1.2 and 1.3 for *FCSNet*, and Supplementary Tables 1.4 and 1.5 for *ImFCSNet*.

**Algorithm 1** 2D or 3D simulation of free diffusing particle

**Input:** $N_{sim}$, $A_{sim}$, $\sigma_{particle}$, $\omega_{xy}$, $np_{step}$, $\sigma^2_{ccd}$, $z_f$, $\omega_z$
**Output:** simulated image stack of size ROI pixels, e.g. 3×3, or ACF curves
▷ $N(0,1)$ denotes sampling from a standard Normal distribution. $Poisson(np_{step})$ denotes sampling from a Poisson distribution with mean of $np_{step}$.

**for** each particle $i$ of the $N_{sim}$ particles **do**
  **if** 2D simulation **then**
    initialize the ($x^i$, $y^i$) position randomly in $A_{sim}$
  **else if** 3D simulation **then**
    initialize the ($x^i$, $y^i$, $z^i$) position randomly in the simulation volume, i.e. $A_{sim} \times 20 \times \omega_z$
  **end if**
**end for**
**for** each frame **do**
  **for** each step in the steps per frame **do**
    **for** each particle $i$ of the $N_{sim}$ particles **do**
      $x^i \leftarrow x^i + \sigma_{particle} \times N(0,1)$
      $y^i \leftarrow y^i + \sigma_{particle} \times N(0,1)$
      **if** 3D simulation **then**
        $z^i \leftarrow z^i + \sigma_{particle} \times N(0,1)$
      **end if**
      **if** the particle moves out of the (2D) simulation area or (3D) volume **then**
        place the particle randomly on the boundary of the (2D) simulation area or (3D) volume
      **end if**
      get the number of photons emitted, $N_{photon} \leftarrow Poisson(np_{step})$
      **if** 3D simulation **then**
        $N_{photon} \leftarrow \lfloor N_{photon} \times \exp\left(-\frac{1}{2}\left(\frac{z}{\omega_z}\right)^2\right)\rfloor$
      **end if**
      **if** 2D simulation **then**
        $\tilde{\omega}_{xy} \leftarrow \omega_{xy}$
      **else if** 3D simulation **then**
        $\tilde{\omega}_{xy} \leftarrow \omega_{xy} + |z| \times z_f$
      **end if**
      **for** each photon of the $N_{photon}$ photons emitted **do**
        $x_{photon} \leftarrow x^i + \tilde{\omega}_{xy} \times N(0,1)$
        $y_{photon} \leftarrow y^i + \tilde{\omega}_{xy} \times N(0,1)$
        **if** $(x_{photon}, y_{photon})$ lies within the ROI **then**
          increase the photon count of the corresponding pixel in the ROI in the current frame
        **end if**
      **end for**
    **end for**
  **end for**
**end for**
**for** each pixel in the ROI **do**
  add the camera noise by sampling from a Gaussian distribution with variance $\sigma^2_{ccd}$ or an empirical EMCCD distribution for each pixel in the ROI
**end for**
**if** we are generating the ACFs **then**
  compute the ACF for each pixel in the ROI and save the ACFs for *FCSNet* training
**else**
  use the simulated ROI image stack for *ImFCSNet* training
**end if**

**Supplementary Table 1.2 :** *FCSNet* simulation range

| Physical parameters | 2D – Setting 1 | 2D – Setting 2 | 3D |
|---|---|---|---|
| $D$ ($\mu m^2/s$) | 0.02 to 15.0 | 0.02 to 15.0 | 0.02 to 15.0 |
| $\omega_{xy0}$ | 0.75 to 0.85 | 0.96 to 1.06 | 1.05 to 1.15 |
| *cps* (thousands) | 4 to 9 | 4 to 9 | 4 to 9 |
| $N_{sim}$ | 358 to 2,389 | 358 to 2,389 | 358 to 2,389 |
| $A_{sim}$ (pixels) | $47 \times 47$ | $47 \times 47$ | $47 \times 47$ |
| ROI (pixels) | $3 \times 3$ | $3 \times 3$ | $3 \times 3$ |
| number of frames | 50,000 | 50,000 | 50,000 |
| steps per frame | 10 | 10 | 10 |

*FCSNet*: Auxiliary physical parameters that we will need for simulation.

**Supplementary Table 1.3 :** *FCSNet* dimensionless simulation parameters

| Simulation parameters | Scale | 2D – Setting 1 | 2D – Setting 2 | 3D |
|---|---|---|---|---|
| $np_{step}$ | natural | 0.424 to 0.954 | 0.824 to 1.854 | 0.816 to 1.836 |
| $\rho$ | log | 0.1623 to 1.0817 | 0.1623 to 1.0817 | 0.1623 to 1.0817 |
| $\sigma_{particle}$ | log | 0.0086 to 0.2350 | 0.0120 to 0.3276 | 0.0071 to 0.19558 |
| $\omega_{xy}$ | natural | 0.6282 to 0.7120 | 0.6805 to 0.7514 | 0.6759 to 0.7403 |
| $\sigma^2_{ccd}$ | natural | 1 to 9 | 1 to 9 | 1 to 9 |

*FCSNet*: the range of dimensionless parameters in our simulation. They are derived from the physical parameters in Supplementary Table 1.1 and Supplementary Table 1.2.

**Supplementary Table 1.4 :** *ImFCSNet* simulation range

| Physical parameters | 2D – Setting 1 | 2D – Setting 2 | 3D – low N & CPS | 3D – high N & CPS |
|---|---|---|---|---|
| $D$ ($\mu m^2/s$) | 0.02 to 50.0 | 0.02 to 50.0 | 0.02 to 50.0 | 0.02 to 50.0 |
| $\omega_{xy0}$ | 0.75 to 0.85 | 0.96 to 1.06 | 1.05 to 1.15 | 1.05 to 1.15 |
| *cps* (thousands) | 1 to 10 | 1 to 10 | 1 to 10 | 5 to 15 |
| $N_{sim}$ | 24 to 486 | 24 to 486 | 24 to 486 | 121 to 608 |
| $A_{sim}$ (pixels) | $15 \times 15$ | $15 \times 15$ | $15 \times 15$ | $15 \times 15$ |
| ROI (pixels) | $3 \times 3$ | $3 \times 3$ | $3 \times 3$ | $3 \times 3$ |
| number of frames | 2,500 | 2,500 | 2,500 | 2,500 |
| steps per frame | 10 | 10 | 10 | 10 |

*ImFCSNet*: Auxiliary physical parameters that we will need for simulation.

We use a larger simulation area of $47 \times 47$ in Supplementary Table 1.2 in contrast to $15 \times 15$ in Supplementary Table 1.4 to account for any potential boundary effect when we simulate large number of frames. We derive the dimensionless simulation parameters based on equation (7). Supplementary Tables 1.3 and 1.5 summarize the corresponding range of simulation parameters. We sample these simulation parameters uniformly during training data generation. Since the values of $\sigma_{particle}$ and $\rho$ cover a wide range, we sample them in a log scale, as indicated under the *Scale* column of these two tables. To illustrate the process, let's denote *mod* and *modSig* as the middle value and the variation of a simulation parameter. In the natural scale, we sample $U[-modSig, modSig]$ and thus the parameter range is $mod - modSig$ to $mod + modSig$. In the log scale, we sample $U[-\log(modSig), \log(modSig)]$ and thus the parameter range is $\exp(\log(mod) - \log(modSig))$ to $\exp(\log(mod) + \log(modSig))$. For example, $\rho$ in Supplementary Table 1.5 is in log scale, and under the column 2D – Setting 1, the range is 0.1082 to 2.1634. It translates to *mod* of 0.4838 and *modSig* of 4.4715.

**Supplementary Table 1.5 :** *ImFCSNet* dimensionless simulation parameters

| Simulation parameters | Scale | 2D – Setting 1 | 2D – Setting 2 | 3D – low N & CPS | 3D – high N & CPS |
|---|---|---|---|---|---|
| $np_{step}$ | natural | 0.106 to 1.060 | 0.206 to 2.060 | 0.204 to 2.040 | 1.020 to 3.060 |
| $\rho$ | log | 0.1082 to 2.1634 | 0.1082 to 2.1634 | 0.1082 to 2.1634 | 0.5408 to 2.7042 |
| $\sigma_{particle}$ | log | 0.0086 to 0.4290 | 0.0120 to 0.5980 | 0.0071 to 0.3571 | 0.0071 to 0.3571 |
| $\omega_{xy}$ | natural | 0.6282 to 0.7120 | 0.6805 to 0.7514 | 0.6759 to 0.7403 | 0.6759 to 0.7403 |
| $\sigma^2_{ccd}$ | natural | 1 to 9 | 1 to 9 | 1 to 9 | 1 to 9 |

*ImFCSNet*: the range of dimensionless parameters in our simulation. They are derived from the physical parameters in Supplementary Table 1.1 and Supplementary Table 1.4.

The respective ranges are in Supplementary Tables 1.3 and 1.5. Lastly, our models predict the $\sigma_{particle}$ in log scale. We can recover the $D$ in unit $\mu m^2/s$ from equation (7) by taking the exponent.

#### 1.2 FCSNet

The simulated training data are ACF curves of 72 points is based on the semi-logarithmic correlator scheme[56]. Briefly, the lag times for which the correlations are calculated first increases linearly $P$ times (e.g., 0 ms, 1 ms, 2 ms ... $P$ ms), then the time step of the increase is doubled and the lag time is increased $P/2$ times ($P$ + 2 ms, $P$ + 4 ms ... ), then the time step is doubled again. The doubling process happens $Q$ - 1 times. So overall the scheme has $Q$ groups in which the time steps are linearly increased, each with double the time step increase as the group before. And the first group contains $P$ correlation values and all subsequent groups contain. In that scheme $P/2$ correlation values. We describe this as a $(P, Q)$ correlator scheme. E.g. for $(P, Q) = (16, 8)$, this results in 16 + 7×8 = 72 values. More information of correlators $P$ and $Q$ are available in the Imaging FCS ImageJ user manual. We excluded $G(\tau = 0)$ from the dataset. We used a mixture of camera noise to improve the robustness of the models. When we simulate the image stacks, we sample from an EMCCD noise data distribution for half of the simulated image stacks and sample from a Gaussian distribution for the other half. The EMCCD noise data is skewed and has a distribution with a long right tail. We varied the variance of the Gaussian noise for each corresponding image stack, and $\sigma^2_{ccd}$ is sampled from $U[1, 9]$. We compute the ACF curves from the simulated image stacks, which we save for training. We performed Z-score standardization on the ACF curves, i.e. subtracting the mean and dividing by the standard deviation of the respective curve.

The total ACF curves we generate for each *FCSNet* training is 479,232. However, we need to filter the ACF curves to remove any ACF with negative $G(\tau)$ values for the first lag. This simple exclusion sifts out ACF curves that can be “very noisy” and may adversely affect the training process. We train the model for 3,000 epochs. We shuffle the data in each epoch, and we group the data into 352 batches, with a batch size of 1,152. As a result, we do not use all of the simulated ACF curves in an epoch, and it introduces some variation in the training data between epochs. Simulation of training data takes about 10.5 hours for 2D free diffusion (and about 12.5 hours for 3D free diffusion) using a GPU. It takes about 12.5 hours to train one network. We use Adam optimizer and an initial learning rate of $10^{-3}$, which we decay exponentially to $10^{-5}$ by the last epoch, i.e. a decay rate ≈ 0.99847 per epoch. We track the training loss and save the model with the lowest training loss as our final model.

#### 1.3 ImFCSNet

The training image stacks of *ImFCSNet* are generated on-the-fly. Each image stack is of the size 2,500 frames × 3×3 pixels. Due to the larger size of the image stacks compared to than ACF curves, we did not save the simulated data or pre-generate the training data. Every image stack is theoretically unique. The *ImFCSNet* training is essentially a single epoch training since we do not re-use the training data. There are

48,000 batches in this single epoch and a batch size of 128 image stacks. We apply Z-score standardization on each image stack.

We use a single GPU for both simulating the training data and training of the model. The training process of *ImFCSNet* is more tedious compared to *FCSNet*. We increase the noise level of our simulated data progressively. We retrain a model multiple rounds, i.e. completing the single epoch every round. We start from Gaussian noise with low variance and increase the variance in each round. We also switch to EMCCD noise towards the end of training. More importantly, we multiply a random factor to the sampled EMCCD noise for every image stack. This random factor is sampled from a uniform distribution (see the last two rows in Supplementary Table 1.6). It scales the EMCCD noise, i.e. if the factor is small, it is less noisy. Based on our observation, using a "full" EMCCD noise, i.e. without multiplying a factor, results in a high training loss that is hard to decrease and the model may not perform well during validation. While the current progressive training schedule is time-consuming, this approach produces trained models that perform more consistently. Perhaps we need many updates to the model to improve its performance[57]. The idea of progressive training is similar to curriculum training[58] where we start with "easy" examples, i.e. low noise level. It is also similar in another context[53], where the authors repeated their training multiple rounds.

In each round, the initial learning rate is reset and the level of data augmentation is increased. We apply an exponential learning rate decay, i.e. starting with the initial learning rate and ending with learning rate, and it's decreased every 32 batches. For example, if the initial learning rate is $10^{-3}$ and final learning rate is $10^{-5}$, then the learning rate decreases by ≈0.99693 every 32 batches. We effectively partition the 48,000 batches (i.e. 1,500 × 32) into 1,500 partitions. Supplementary Table 1.6 summarizes our training schedule. Speeding up the training process of *ImFCSNet* is certainly an area for improvement. Due to the longer training schedule, it takes about 130 and 180 hours for 2D and 3D settings, respectively, to train one network. Improving the training speed is one area of improvement.

**Supplementary Table 1.6 :** Training schedule of *ImFCSNet*

| Round | noise type | variance / factor | initial learning rate | final learning rate |
|---|---|---|---|---|
| 1 | Gaussian | 0.09 to 0.11 | $10^{-3}$ | $10^{-5}$ |
| 2 | Gaussian | 0.1 to 1.9 | $10^{-3}$ | $10^{-5}$ |
| 3 | Gaussian | 1.0 to 5.0 | $10^{-3}$ | $10^{-5}$ |
| 4 | Gaussian | 1.0 to 9.0 | $10^{-3}$ | $10^{-5}$ |
| 5 | Gaussian | 1.0 to 9.0 | $10^{-3}$ | $10^{-5}$ |
| 6 | Gaussian | 1.0 to 9.0 | $10^{-3}$ | $10^{-7}$ |
| 7 | EMCCD | 0.1 to 0.5 | $10^{-3}$ | $10^{-7}$ |
| 8 | EMCCD | 0.1 to 1.0 | $10^{-3}$ | $10^{-7}$ |

We reload a previously trained model from round 2 onward and change the random number generator seed in each round of training. We sample the variance of the Gaussian noise or the multiplicative factor, i.e. $EMCCDnoisefactor$, to the EMCCD noise from a uniform distribution for each simulated image stack. The range of the uniform sampling is in the column *variance/factor*. We use Adam optimizer with an exponential learning rate decay, with the starting and ending learning rates as shown.

We change the random number generator seed each round to ensure we have different simulated data. Like *FCSNet*, we track the training loss, and the model with the lowest training loss is the "best model" for each round. From round 2 onward, we reload the best model of the previous round and restart the training. In addition, we reload the model with the lowest training loss from rounds 4 and 5 in round 6. In other words, we reload the model from round 5 if the model has a lower training loss than the counterpart from round 4; otherwise, we retrain the model from round 4. Similarly, when we are at round 7, we will reload the best model from rounds 4 to 6 for further retraining. Supplementary Table 1.6 describes the settings we used for the 2D and 3D training. Finally, we use the model from the last round for evaluations.

### 2 Evaluation of simulated test data

We chose 4 diffusion coefficients: 0.1, 1, 5, and 10 $\mu m^2/s$. We generated 50 image stacks for each $D$ by randomizing the CPS, the number of particles in the simulation area, and the seed number of the random number generator. There are 50,000 frames in each image stack. For each simulated test image stack, we added either Gaussian noise and varied the variance by sampling the variance uniformly from $U[1, 9]$, or sampled from the empirical EMCCD noise and multiplied by a factor from $U[0.1, 1.0]$, as mentioned in Supplementary Section 1.3. The size of our test data is thus 400 (i.e. 4 $D$'s $\times$ 50 image stacks $\times$ 2 types of noise). Since there is no bleaching in the simulated test data, we do not apply polynomial bleach correction to the dataset.

We compared our results against NLS 1×1 evaluations with 50,000 frames, and in the case of *ImFCSNet*, we included the evaluations of NLS with 2,500 frames. The results in Supplementary Fig. 2.1 show that the predictions by the models are centered around the ground truths for most of the test data. The values match the NLS results as the scatter plot falls along the diagonals. We excluded data points where NLS predicts $\leq 100\mu m^2/s$. In total, we have remaining 379 and 396 test data points for 2D and 3D evaluations, respectively. By comparing the difference between the log of the predicted $D$ and the log of the actual $D$, the means and standard deviations are 1) in the 2D case, NLS 50,000 frames: 0.050 $\pm$ 0.059, NLS 2,500 frames: 1.223 $\pm$ 0.529, *FCSNet*: 0.020 $\pm$ 0.126, *ImFCSNet*: 0.022 $\pm$ 0.152, and 2) in the 3D case: NLS 50,000 frames: 0.093 $\pm$ 0.261, NLS 2,500 frames: 1.289 $\pm$ 1.384, *FCSNet*: -0.016 $\pm$ 0.046, *ImFCSNet*: 0.041 $\pm$ 0.077. The outliers of *FCSNet* and *ImFCSNet* predictions, in particular in Supplementary Fig. 2.1c, contribute to the higher standard deviations in the log scale for the 2D case. The differences from ground truth in natural scale are: 1) in the 2D case, NLS 50,000 frames: 0.207 $\pm$ 0.486, NLS 2,500 frames: 1.346 $\pm$ 2.934, *FCSNet*: -0.001 $\pm$ 0.297, *ImFCSNet*: -0.027 $\pm$ 0.231, and 2) in the 3D case, NLS 50,000 frames: 0.596 $\pm$ 2.769, NLS 2,500 frames: 7.138 $\pm$ 17.451, *FCSNet*: -0.148 $\pm$ 0.201, *ImFCSNet*: 0.067 $\pm$ 0.257. Importantly, the results demonstrate that NLS evaluations with 2,500 frames are biased and fail to recover the diffusion coefficients. We depict the CNN results in Supplementary Fig. 2.1 with different colours for the different noise type. We observe instances of higher prediction errors by CNNs (and also NLS) when using the EMCCD noise distribution, i.e. a more noisy background.

### 3 Reproducible training

We trained four different models and compared them against each other to demonstrate reproducible training of *FCSNet* and *ImFCSNet*. For that purpose we used 1) the DOPC lipid bilayer in Figs. 2a,d and 2) the in-focus 100 nm bead solution measurements in Figs. 2b,c. We use different seeds for the training of different models, to change the simulated data and their sampling, and randomly initialize the model weights. In Supplementary Fig. 3.1, we observe that the pairwise comparisons of the different models are distributed around the diagonals, indicating that the predictions are close. There are some variations, as anticipated, since the weights of the trained models are different. Nonetheless, the models seem to provide similar predictions. The mean and standard deviation along the ($x$, $y$) axes for the pairwise comparison are, notably: Supplementary Fig. 3.1a: (0.817 $\pm$ 0.032, 0.818 $\pm$ 0.032), Supplementary Fig. 3.1b: (1.258 $\pm$ 0.032, 1.265 $\pm$ 0.032), and Supplementary Fig. 3.1e: (1.234 $\pm$ 0.029, 1.234 $\pm$ 0.028). The smaller standard deviations point to higher precision of the different trained models. For Supplementary Fig. 3.1d, the corresponding values are (0.721 $\pm$ 0.105, 0.725 $\pm$ 0.105). Although the standard deviation is larger, the mean of $x$ and $y$ values are virtually on the diagonal.

In Supplementary Fig. 3.1c, the *ImFCSNet* models are trained with low SNR simulated 3D diffusion data, i.e. with lower $N$ and $CPS$ as indicated in Supplementary Table 1.5. The pairwise comparison of the models shows more pronounced variation visually. The corresponding mean and standard deviation along the ($x$, $y$) axes are (1.268 $\pm$ 0.106, 1.319 $\pm$ 0.115). There is a greater difference in the mean values, as well as, the standard deviations. Put differently, the differences in the outputs of the four *ImFCSNet* models are larger. Intuitively, it is harder to train the networks when the training data is noisy. In contrast, we see an improved comparison when the models are trained with high SNR training data, as seen in Supplementary Fig. 3.1e. In

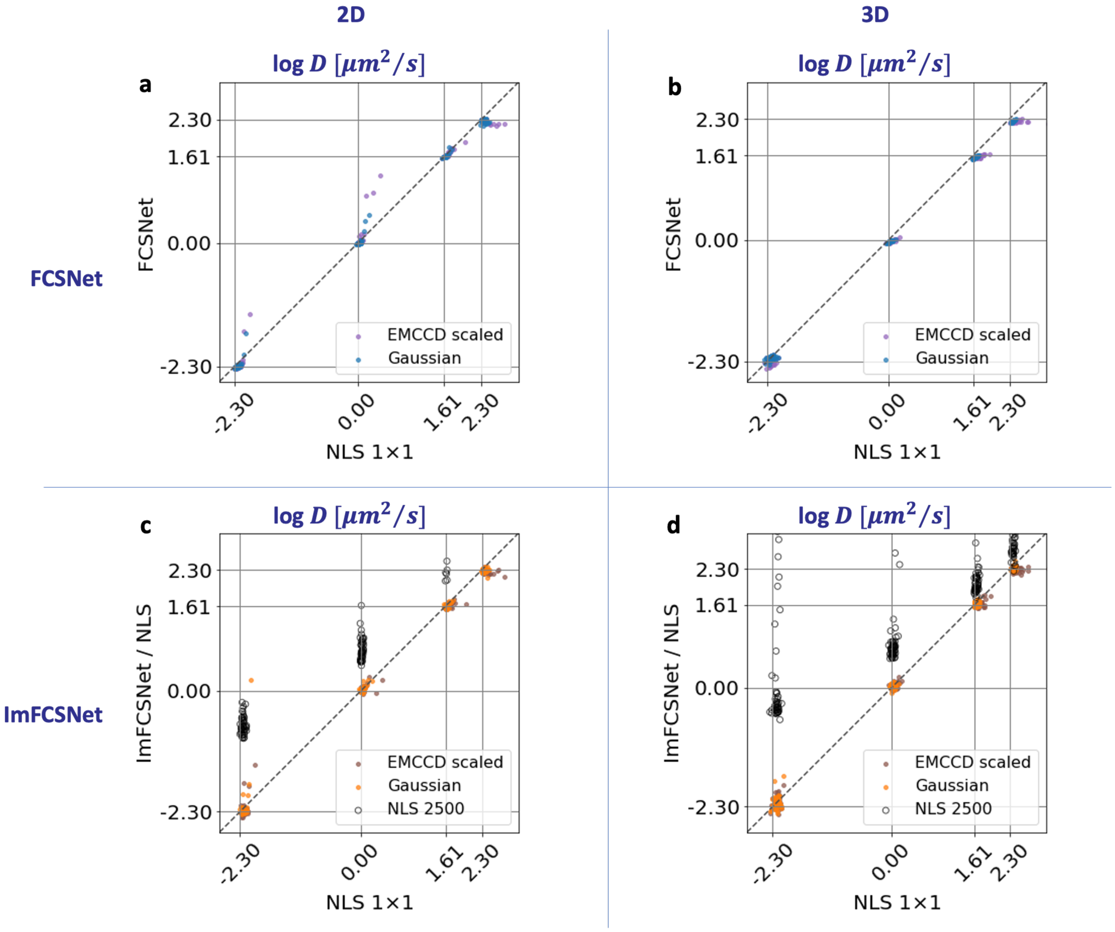


**Supplementary Figure 2.1 : Simulated test data**. The plots are in log scale. The ground truth diffusion coefficients of the test data are 0.1, 1, 5, and 10 $\mu m^2/s$. **a**, **b**, *FCSNet* and **c**, **d**, *ImFCSNet* models can recover the ground truth for most image stacks and the performances are comparable to NLS evaluations. In the lower row, we observe a positive bias in the predictions by NLS for 2,500 frames.

Supplementary Fig. 3.1f, we observe that the model trained with high SNR seems to become more sensitive to out-of-focus measurements, as we can see a steeper slope in the predictions (in green) when the sample de-focused. There is a benefit in training *ImFCSNet* with low SNR training data, but the price to pay is more training. The result here also underlines the importance of training multiple models and calibrating against NLS predictions, as shown in Fig. 2, to choose the best model for eventual implementation.

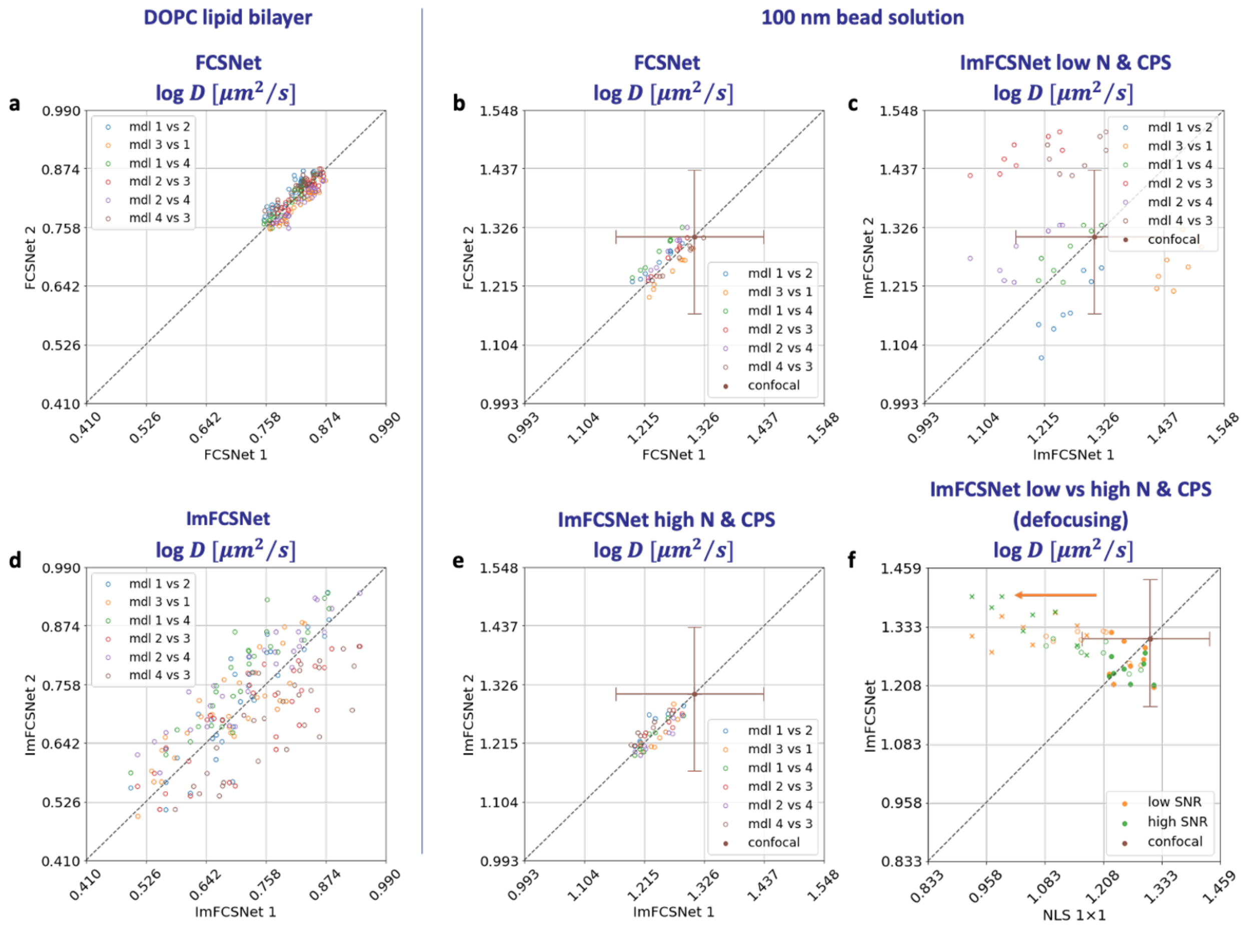


**Supplementary Figure 3.1 : Reproducible performances of randomly initialized models, trained with different simulated data**. We train 4 *FCSNet* and *ImFCSNet* models, respectively. The pairwise comparisons of the predictions on the 1) DOPC lipid bilayer and 2) the 100 nm bead solution in **a** – **e** show how close the predictions are to each other. The results indicate that the performances are reproducible. The different models are randomly initialized and trained with different datasets. However, the performances on different *ImFCSNet* in **c** are inconsistent. We can improve the precision by training the model with a higher signal dataset, i.e. by increasing N and CPS, as shown in **e**. **f**, albeit *ImFCSNet* obtained with high SNR training data still perform well against out-of-focus data, the green plot shows a slight increase in $D$ in the direction of defocusing.

### 4 Experimental measurements

#### 4.1 Preparation of DOPC and POPC supported lipid bilayer

Supported Lipid Bilayers (SLBs) were created using either 1,2-dioleoyl-sn-glycero-3-phosphocholine (DOPC) or 1-palmitoyl-2-oleoyl-sn-glycero-3-phosphocholine (POPC). Both DOPC and POPC are fluid at room temperature. SLBs were prepared using the optimized vesicle fusion protocol[47]. A calculated amount of lipid and lipid dye solution (0.01 mol% 14:0 Liss Rhod PE) in chloroform were mixed well into a round bottom flask. This was followed by solvent evaporation for 2-3 h in a rotary evaporator (Rotavap R-210, Buchi, Switzerland). This lipid film obtained was re-suspended in 2 mL buffer containing 10 mM HEPES and 150 mM NaCl (pH 7.4). Small unilamellar vesicle (SUV) were prepared by sonicating the suspension until clarity using bath sonicator (Elmasonic S30H, Elma Schmidbauer GmbH, Singen, Germany). 200$\mu$L of SUV solution was added to 200$\mu$L of the same buffer into an O-ring (1.5 cm inner diameter prepared from silicon elastomer SYLGARD 184 Silicone Elastomer Kit, Dow, Michigan, USA) attached to a glass cover slide (24×50-1, Fisher Brand Microscope, Thermo Fisher Scientific), incubated above lipid melting point at 65 °C for 1 h to allow vesicle fusion and formation of SLB. The sample was slowly cooled to room temperature (25 °C) for 1 h before removing the unfused vesicle by washing with the same buffer. SLB measurements were performed at 25 °C for DOPC and 37 °C for POPC.

#### 4.2 Cell culture and transfection

**Plasmid:** GPI-GFP plasmid having a glycosylphosphatidylinositol-anchored protein tagged with a green fluorescent protein was a kind gift from John Dangerfield (Anovasia Pte Ltd, Singapore). The construction of the PMT-mEGFP plasmid consisting of a plasma membrane targeting sequence fused with mEGFP tag has been described in a previous publication[28]. A PMT-mEGFP plasmid was digested with NheI (NheI-HF, #R3131S, New England Biolabs, Massachusetts, USA) and HindIII. The digested PMT-mEGFP PCR products were then ligated (SpeI and NheI are isocaudomers) using T4 DNA ligase (#M0202S, New England Biolabs, Massachusetts, USA).

**Plating:** Adherent HeLa or CHO-K1 cells were cultivated in DMEM (Dulbecco's Modified Eagle Medium (Invitrogen, Carlsbad, CA) supplemented with 10 % fetal bovine serum (Invitrogen, Carlsbad, CA) and 1 % PS (penicillin and streptomycin) at 37 °C in 5 % $CO_2$. Once the culture is approximately 90% confluent $1 \times 10^4$ cells were seeded on the glass-covered dishes (35 mm No. 1.0 cover glass $0.13 - 0.16$ mm, MatTek, Massachusetts, USA) supplemented with DMEM + 10% FBS for $\approx$ 24 h followed by transfection.

**Transfection:** CHO-K1 (CCL-61™, ATCC, Manassas, Virginia, USA) or HeLa cells were transfected with either PMT-mEGFP or GPI-GFP, respectively. Transfected cells were washed with HBSS (Hank's Balanced Slat Solution) (Gibco, Thermo Fisher Scientific, Massachusetts, USA) once and imaged under TIRFM with Phenol red free DMEM + 10% FBS at 37 °C.

Instructions on cell culture and transfection for preparation of live-cell samples can be found on Protocol Exchange[59].

#### 4.3 Fluorescent beads preparation

100 nm TetraSpeck beads (Invitrogen, Carlsbad, CA) were diluted by a factor of 1 to 100 for SPIM-FCS and confocal FCS respectively followed by sonication to minimize aggregation. For SPIM-FCS, liquid sample was either mounted in a transparent, heat-sealed bag made from fluorinated poly-ethylene-propylene (FEP) films with refractive index of 1.341-1.347, which is close to that of water. For confocal FCS, a drop of bead solution was placed on top of coverslip and measured at 30 $\mu$m above to exclude non-specific binding between the glass surface and the beads.

#### 4.4 Drosophila embryo measurement

The eGFP-bicoid transgenic fly line used in this work was a gift from Thomas Gregor's lab[29]. The eGFP::Bcd line used here was created for another work done[36]. eGFP::Bcd was crossed with His2A::mCherry to label the nuclei of early blastoderm embryos.

eGFP::Bcd; His2A::mCherry embryos at n.c. 9 were dechorionated and imaged at n.c. 14. These embryos were mounted in a notch opening created in a square FEP tube (2 mm × 2 mm; Adtech Polymer Engineering, England, United Kingdom) and embedded in low melting 1% conc. agarose(UltraPureTM Low Melting Point Agarose, 16520100, Thermofisher Scientific, United States) in water. This open mounting approach was adopted so that the light sheet struck the immobilised embryo, and the fluorescence emission could be captured without any scattering or signal losses from the thick FEP tube. The embryo was positioned so that the anterior-posterior axis coincided with the plane of the light sheet, and the detection objective imaged the embryo's coronal or sagittal plane.

#### 4.5 Instrumentation and data acquisition

**TIRFM:** Imaging total internal reflection FCS (ITIR-FCS) was performed with two objective-type TIRFM systems. The first system is an Olympus microscope (IX83) with a motorized TIRF illumination combiner (cellTIRF-4Line IX3-MITICO, Olympus), with an oil-immersion objective (Apo N, 100×, NA 1.49, Olympus) equipped with 488 nm and 561 nm. Second system is total internal reflection fluorescence microscope (IX-71, Olympus, Japan) with an oil-immersion objective (PlanApo, 100×, NA 1.45, Olympus) coupled with TIRF illuminator model IX2-RFAEVA-2 (Olympus), equipped with 488 nm LuxX+ 150 mW (Omicron-Laserage Laserprodukte GmbH) and 532 nm LS 150 mW (OBIS, Coherent, CA) lasers were combined with a Omicron LightHUB+ laser combiner (Omicron-Laserage Laserprodukte GmbH) into a single fibre output before focusing light to the back focal plane of the TIRF objective. For detection purposes, we employed an iXon DU860 (Andor, Oxford Instrument, UK) electron-multiplying charge-coupled device (EMCCD) with a pixel size of 24 $\mu$m. This was controlled through a custom-built data acquisition software. We adjusted the focus in real-time by maximizing the amplitude of the ACFs[24]. The camera was operated at - 80 °C with the following settings: Baseline clamp = "on", pixel readout speed = 10 MHz, maximum analog-to-digital gain = 4.7, vertical shift speed = 0.45 $\mu$s, electron-multiplying gain = 300.

**Confocal microscope:** For the sole purpose of comparing diffusion measurements of fluorescence beads in water, an Olympus FV 1200 laser scanning confocal microscope (IX83, Olympus, Japan) was employed, equipped with a PicoQuant time-resolved LSM upgrade kit (Microtime 200, GmbH, Germany). We illuminated the fluorescence bead sample with a pulsed 485 nm laser(LDH-D-C-488, PicoQuant), operating at a repetition rate of 20 MHz. This was reflected in the back focal plane of an Olympus UPLSAPO 60X/1.2 NA water immersion objective. The emitted signal passes through a 120 $\mu$m pinhole before being filtered by a 510/23 emission filter (Semrock, US) before detection by a single photon sensitive avalanche photodiodes (SAPD) (SPCM-AQR-14, PerkinElmer). The captured signal was then analyzed using SymPhoTime 64 (PicoQuant, Germany) to compute for ACFs free from background related-artefact, which were subsequently fitted with a 3D free diffusion single-particle model[40].

**SPIM:** The drosophila embryos were imaged using a home build SPIM setup previously described[3,25]. The illumination section consists of a 488 nm diode laser (Cobolt 06-MLD 488nm 0488-06-01-0100-100, Cobolt AB, Sweden)and a 561 nm diode laser(Cobolt 06-DPL 561nm 0561-06-91-0100-100, Cobolt AB, Sweden). The 488 nm laser was used to illuminate the eGFP bicoid, and the 561 nm laser was used to illuminate the His2A:: mCherry to locate the nuclei.

The two lasers were combined and directed through an optical fibre (kineFLEXP-3-S-405..640-1.0-4.0-P2, Qioptiq, United States). The laser beams were further expanded two fold by an off-axis parabolic mirror pair (reflected focal lengths of 2 inches: MPD129-G01 and 4 inches: MPD149-G01, Thorlabs Inc., United States) based beam expansion system so that the beam filled the back aperture of the illumination objective. After the beam expansion, the beam was passed through an achromatic cylindrical lens of 75 mm focal length (ACY254-075-A; Thorlabs Inc., United States) and an illumination objective (SLMPLN 20 ×/0.25

NA; Olympus, Japan) to form the light sheet. The light sheet thickness obtained had a $\frac{1}{e^2}$ radius of approximately 1.1 $\mu$m. The thinnest section of the light sheet was aligned to coincide with the focal plane of the detection objective. (LUMPLFLN 60×/1.0 NA, Olympus, Japan).

The mounted embryos were imaged in a custom-made cube chamber filled with water. The cube had dimensions of 3 cm × 3 cm × 3 cm with an opening on the top for mounting the sample and an opening on one side for a mounting hole for placing the water dipping detection objective. The FEP tube containing the mounted embryos was held by self-closing forceps and mounted on a motorised piezoelectric stage, allowing 3-axis movements and rotation (Q-545 Q-MotionR Precision Linear Stage; Physik Instruments, Germany). The embryos were lowered into the sample chamber and moved so as to position the embryo to be illuminated by the light sheet. The detection objective was mounted on a piezo flexure objective scanner(P-721 PIFOC; Physik Instruments, Germany) for fine controlling the detection objective's position with respect to the light sheet.

For FCS-based measurements, the emission obtained by the detection objective was passed through a filter (FF03-525/50-25, Semrock, United States) and was projected onto an EMCCD camera (Andor iXON3, 860, 128 × 128 pixels, Andor Technology, United Kingdom) by a tube lens (LU074700, f=180 mm, Olympus, Tokyo, Japan). Before the FCS measurement, a sCMOS camera (OCRA-Flash 4.0, V2; C11440 Japan) was used to visualise the global position of the embryo to decide the region of interest. The embryos experienced 30 $W/cm^2$ of laser intensity to limit photobleaching. A region on the anterior dorsal side of the embryo with numerous nuclei was chosen at the $14^{\text{th}}$ division cycle and imaged on the EMCCD camera to perform imaging FCS. For imaging FCS measurements, 50,000 frames were recorded with an exposure time of 2 ms (per frame time = 2.04 ms)

### 5 Additional DOPC lipid bilayer plots

Supplementary Figs. 5.1a–c are additional plots to show the consistency in the prediction trend by *FCSNet* and *ImFCSNet* in comparison to NLS over 300,000 frames. In Supplementary Fig. 5.1d, we evaluate the ACF curves and NLS fits of the first 50,000 frames of Supplementary Fig. 5.1a. In Supplementary Fig. 5.1e, we show an example of a fit from Supplementary Fig. 5.1d, where NLS fits the ACF curve well. Furthermore, *FCSNet* predicts a $D$ of 2.40 $\mu m^2/s$, which is in good agreement with the prediction of 2.38 $\mu m^2/s$ by NLS. *ImFCSNet* predicts 2.94 $\mu m^2/s$ and NLS 3×3 predicts 1.78 $\mu m^2/s$ for a 3×3 area encompassing that pixel. On the other hand, Supplementary Fig. 5.1f shows an ACF curve that was not properly approximated by an NLS fit, which predicts $D$ of 0.97 $\mu m^2/s$ and *FCSNet* predicts 1.77 $\mu m^2/s$, while *ImFCSNet* predicts 2.72 $\mu m^2/s$ and NLS 3×3 predicts 1.51 $\mu m^2/s$ (for a 3×3 area encompassing that pixel). The observations are similar in Supplementary Figs. 5.1g–i. This indicates that at least to some extent that *FCSNet* and *ImFCSNet*, are less sensitive to artefacts and can still extract values closer to the expected D compared to NLS fits.

The time resolution of *ImFCSNet* is 20 times better than for an NLS fit. This raises the question whether *ImFCSNet* actually predicts correct trends even at such short evaluation times of 2.5 s. We zoom in to Supplementary Fig. 5.1c to analyze the *ImFCSNet* predictions, in particular the steep decrease in $D$ values in the region of 140 s to 170 s. We compute NLS predictions in a sliding window manner with a step-size of 10,000 frames. Although this leads still to averaging over 50,000 frames, we get predictions now in a step size of 10,000 frames. It is evident that we see the downward trend in these NLS predictions in Supplementary Fig. 5.2 matching the trend of *ImFCSNet* predictions. The NLS $D$ values are higher than *ImFCSNet* because it still computes the average of 50,000 frames. It nonetheless reaches the lowest $D$ value of 1.70 $\mu m^2/s$ at 190 s, which coincides with the time when the majority of the low $D$ values predicted by *ImFCSNet* fall within the preceding 50,000 frames. This observation implies that *ImFCSNet* is capturing the underlying events at a finer time-resolution.

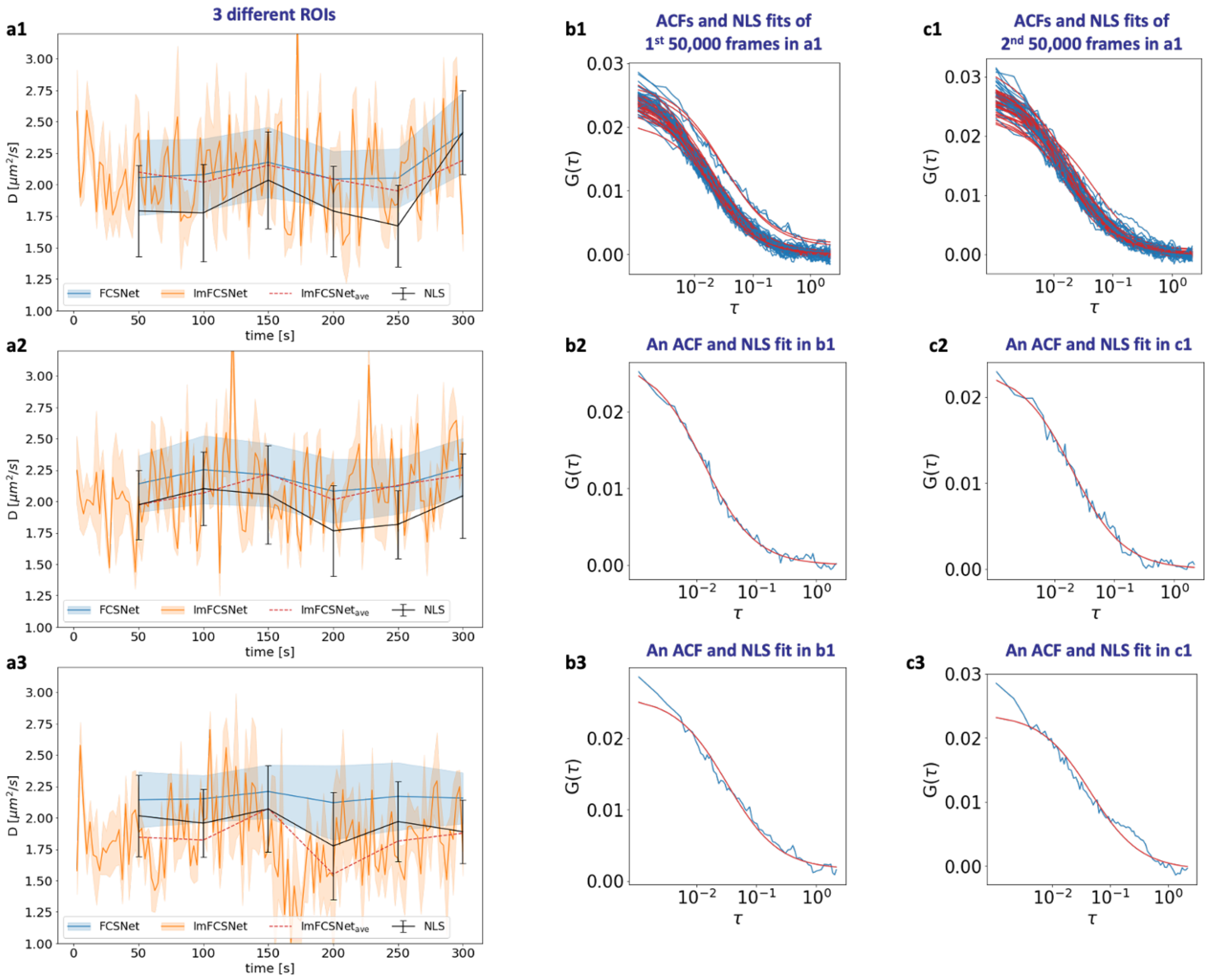


**Supplementary Figure 5.1 : Additional DOPC lipid bilayer analysis**. **a1**,**a2**,**a3**, evaluations of three different 6×6 ROIs of the DOPC lipid bilayer measurements over 300,000 frames. **b1**, ACFs and NLS 1×1 fits of the first 50,000 frames in **a1**. **b2**, an ACF and NLS fit from **b1**. NLS predicts $D$ of 2.38 $\mu m^2/s$, while *FCSNet* predicts 2.40 $\mu m^2/s$ and *ImFCSNet* predicts 2.94 $\mu m^2/s$ for a 3×3 area encompassing that pixel. **b3** is an example of misfit in **b1**. NLS predicts $D$ of 0.97 $\mu m^2/s$ while *FCSNet* predicts 1.77 $\mu m^2/s$ and *ImFCSNet* predicts 2.72 $\mu m^2/s$ (for a 3×3 area encompassing that pixel). **c1**, ACFs and NLS 1×1 fits of the next 50,000 frames (i.e. 50,001$^{th}$ to 100,000 frames in **a1**). **c2**, an ACF and NLS fit from **c1**. NLS predicts 1.59 $\mu m^2/s$, *FCSNet* predicts 1.67 $\mu m^2/s$, and *ImFCSNet* predicts 1.81 $\mu m^2/s$ and NLS 3×3 predicts 1.59 $\mu m^2/s$ (for a 3×3 area encompassing that pixel). **c3** is an example of misfit in **c1**, NLS predicts 0.68 $\mu m^2/s$ and *FCSNet* predicts 1.39 $\mu m^2/s$, while *ImFCSNet* predicts 2.10 $\mu m^2/s$ and NLS 3×3 predicts 1.03 $\mu m^2/s$ (for a 3×3 area encompassing that pixel).

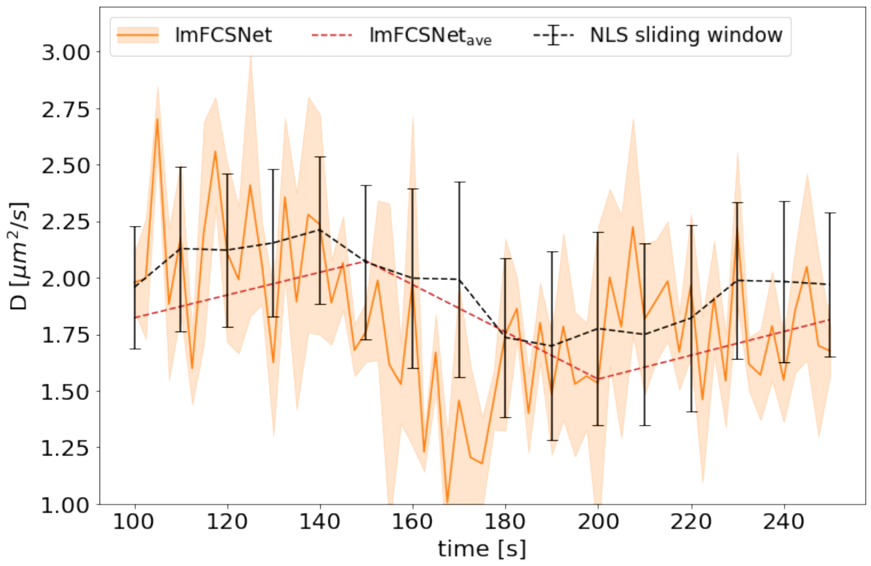


**Supplementary Figure 5.2 : Comparison with NLS 1×1 sliding window of 50,000 frames**. Further to Supplementary Fig. 5.1c, we include the NLS 1×1 sliding window evaluations with a step-size of 10,000 frames for each evaluation, which is shown by the black dotted line. The NLS predictions follow the trend of the *ImFCSNet* predictions. In particular, NLS predictions reaches the lowest $D$ value of 1.70 $\mu m^2/s$ at 190 s when majority of the low $D$ values predicted by *ImFCSNet* fall within the preceding 50,000 frames.

### 6 CHO-K1 cell expressing PMT-mEGFP measurement

Supplementary Fig. 6.1a shows the the whole cell averaged over the first 2,500 frames, of which a 21×21 ROI (indicated in green and shown in Supplementary Fig. 6.1b) is used for evaluations in Fig. 3. Supplementary Fig. 6.1c shows the ACFs for the 441 pixels. Supplementary Figs. 6.1d,e show the N and $D$ maps for the ROI, respectively, as determined by NLS. Supplementary Fig. 6.1e is the same diffusion map in Fig. 3a.

### 7 Additional discussion of drosophila embryo measurements

We averaged the first 2,500 frames of the 50,000 frame stack of a drosophila light sheet image stack to create Supplementary Fig. 7.1a. The heatmap of the average intensities of the first 2,500 frames, 3×3 binning, is in Supplementary Fig. 7.1b. The brighter regions correspond to the nuclei. We superimpose the green dots on the heatmap to depict areas with the predictions by *ImFCSNet* that are larger than 0.1 $\mu m^2/s$. The faster diffusing particles are found between the nuclei as also predicted by NLS. However, the values are mostly lower than those obtained by NLS. In Supplementary Fig. 7.1c, we see the corresponding points of $D$ larger than 0.1 $\mu m^2/s$ in the *ImFCSNet* diffusion map in Fig. 4e. Therefore, *ImFCSNet* makes sensible predictions in terms of the diffusion pattern with areas of higher diffusion being identified and thus producing a $D$ map that is at least partially consistent with expectations. However, the values are mostly predicted at much lower values. So this is a clear case where *ImFCSNet* fails, unlike *FCSNet*. This raises the question why *ImFCSNet* fails. We plot the ACF curves of two 6×6 ROI regions in Supplementary Figs. 7.1d,e, which correspond to regions between nuclei and within a nucleus, respectively. The curves, although containing some artefacts to movement, as mentioned before, still contain a decaying ACF which NLS and *FCSNet* can exploit, thus supporting the consistency between their results. We also find in Supplementary Section 9 that the SNR of the drosophila embryo measurement to be relatively high. These observations suggest that the evaluations of *ImFCSNet* on drosophila measurement is affected by another factor. As *ImFCSNet* does not evaluate the ACF but the changing intensities due to the molecular movement of particles, we hyothesize that it is more sensitive the actual dynamics. With recent publications indicating that bicoid dynamics is

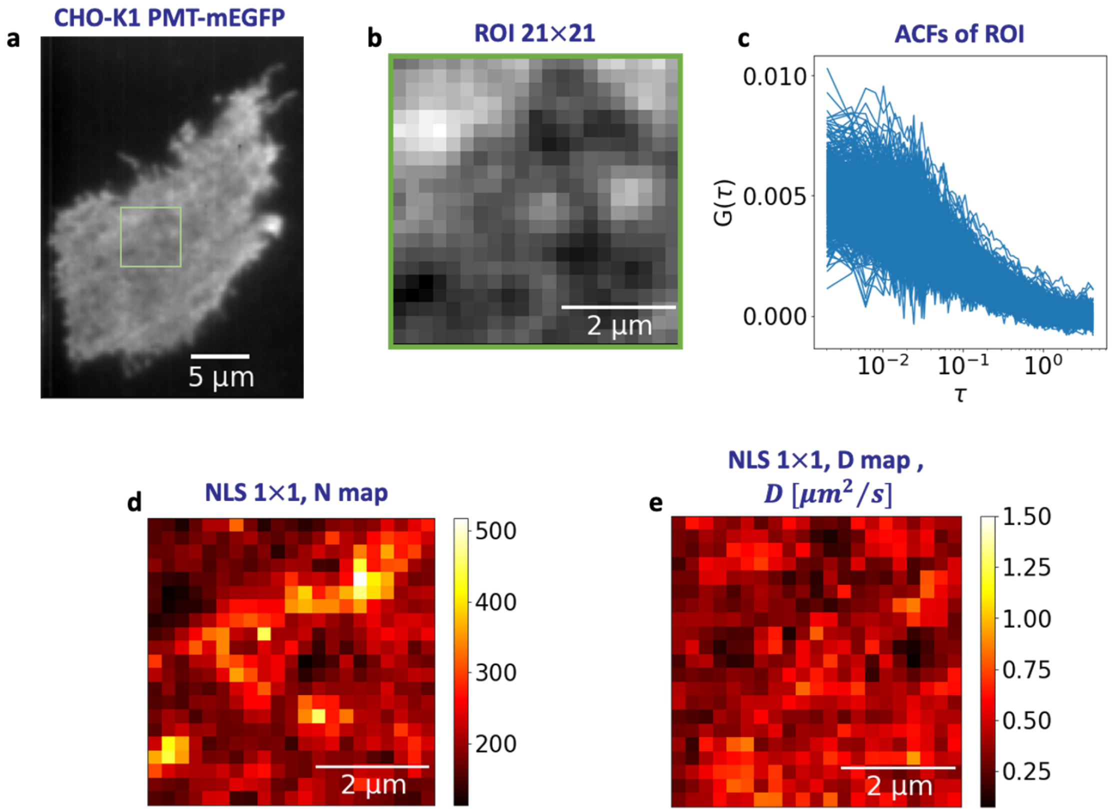


**Supplementary Figure 6.1 : CHO-K1 cell expressing PMT-mEGFP**. **a**, wide-field image of the measurement. **b**, The 21×21 ROI evaluated in Fig. 3. **c**, ACF curves of the ROI. NLS evaluation returns multiple parameters – N map in **d**, and $D$ map in **e**.

better described by a two component diffusion process, we hypothesize that we may need to include multi components into the CNN training for *ImFCSNet* to be able to predict correct values.

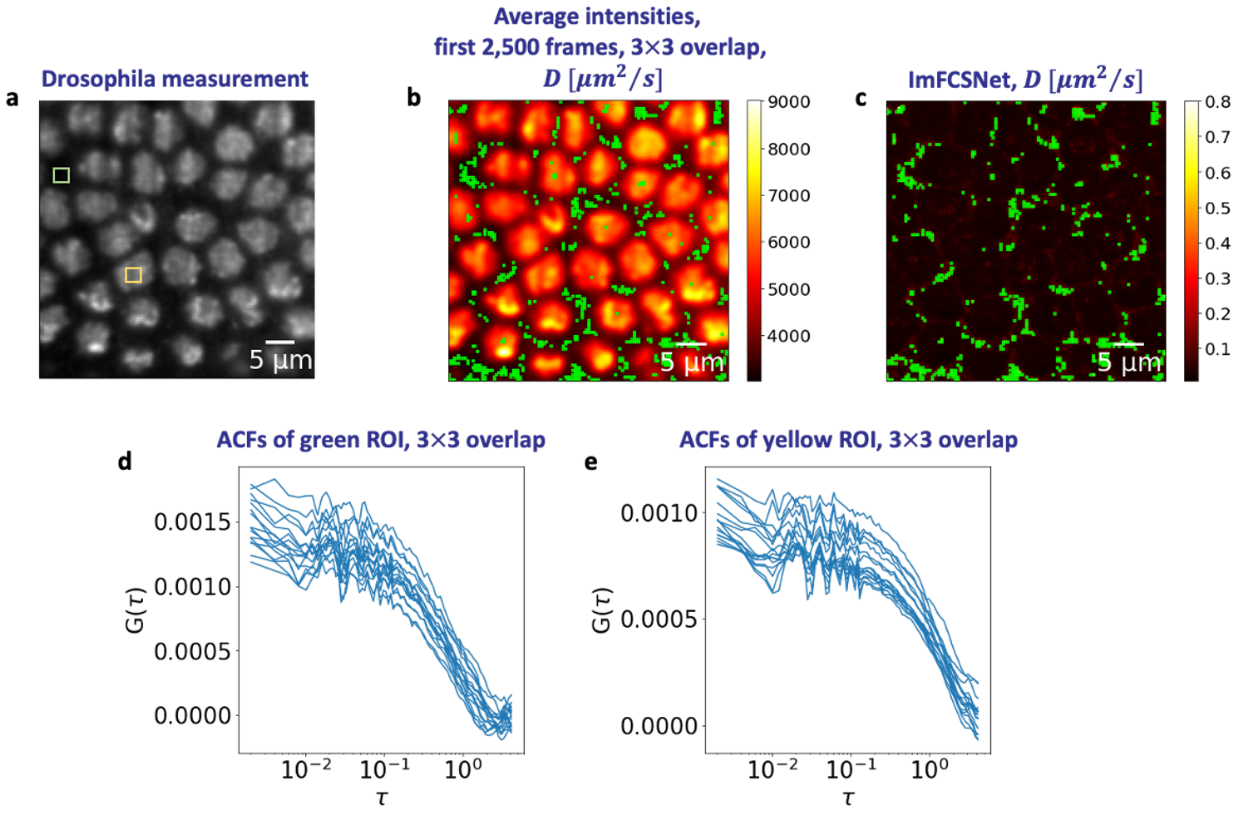


**Supplementary Figure 7.1 : Regions in the drosophila embryo measurement where *ImFCSNet* predicts $D > 0.1\ \mu m^2/s$.** **a**, Wide field image, averaged over the first 2,500 frames. **b**, Heat map of the average intensities, with 3×3 overlapping binning, of the first 2,500 frames. We superimpose the green dots on the heatmaps to indicate the pixels where predictions by *ImFCSNet* are above 1 $\mu m^2/s$. *ImFCSNet* can identify the faster diffusion between the nuclei. The brighter spots correspond to the nuclei, and we expect the diffusion coefficients to be slower. **c**, the corresponding locations with diffusion coefficients are above 0.1 $\mu m^2/s$ on the diffusion map of *ImFCSNet* in Fig. 4e. **d**, ACF curves overlapping 3×3 binning for the 6×6 yellow ROI in **a** for 50,000 frames. **e**, ACF curves of overlapping 3×3 binning for the 6×6 green ROI in **a** for 50,000 frames.

### 8 Varying the laser power of HeLa GPI-GFP measurements

We demonstrate in this section the importance of having sufficient signals in the measurements. Supplementary Fig. 8.1 shows the different *ImFCSNet* results at different laser power. We apply *ImFCSNet* on a series of 4 measurements of 2,500 frames taken at different laser power. We did not use bleach correction due to the low frame count. We also have a measurement of the same HeLa cell at a laser power of 196 $\mu$W (1.80 W/cm$^2$) for 50,000 frames. We evaluate this longer measurement with NLS and use the result as the baseline, which is on the x-axis. In Supplementary Figs. 8.1c,d, we observe a larger variation in *ImFCSNet* predictions for the measurements with a low laser power of 47 $\mu$W (0.43 W/cm$^2$) or 106 $\mu$W (0.97 W/cm$^2$). In Supplementary Figs. 8.1e,f, the predictions improve as the laser power increases to 196 $\mu$W (1.80 W/cm$^2$) and above. At low laser power, the predictions by *ImFCSNet* can become very large.

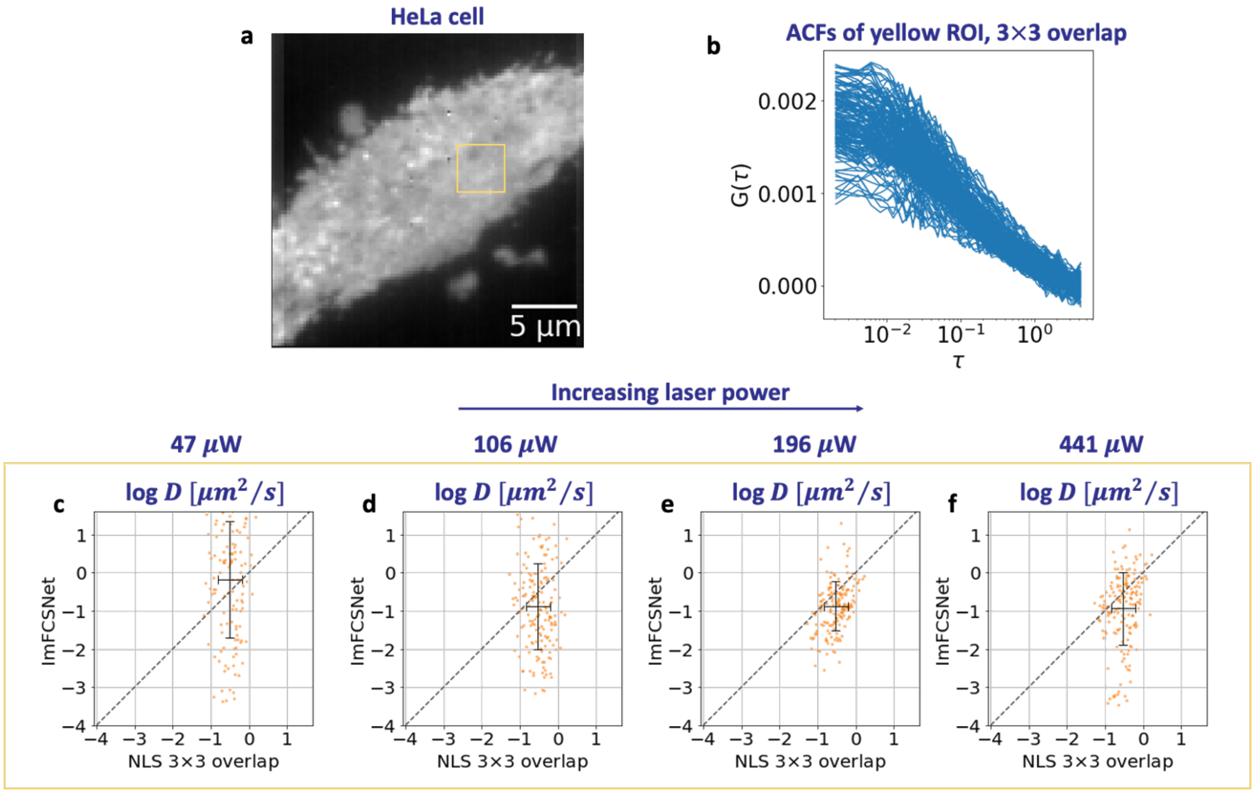


**Supplementary Figure 8.1 : HeLa cell expressing GPI-GFP**. **a**, we evaluate a 15×15 ROI highlighted in yellow on a HeLa GPI-GFP measurement. **b**, the ACF curves computed at 3×3 overlapping binning for the yellow ROI. Different measurements are taken at an increasing laser power, from left to right: 47 $\mu$W (0.43 W/cm$^2$) in **c**, 106 $\mu$W (0.97 W/cm$^2$) in **d**, 196 $\mu$W (1.80 W/cm$^2$) in **e**, and 441 $\mu$W (4.05 W/cm$^2$) in **f**. These measurements contain 2,500 frames. We use the NLS prediction of a 50,000 frame measurement of the same cell at 196 $\mu$W (1.80 W/cm$^2$) as a reference on the x-axis of all the plots. The predictions by *ImFCSNet* vary to a larger degree (on the y-axis) when the laser power is low, and the precision improves as the values lie closer to the diagonal when the laser power increases to 196 $\mu$W (1.80 W/cm$^2$) and above. This example demonstrates that the performance of *ImFCSNet* depends on the strength of the signals in the measurements. The black crosses in the scatter plots are the average and $\pm 1$ standard deviation of the respective log $D$ values. The standard deviation of NLS predictions is 0.31. For *ImFCSNet* predictions, the standard deviations are 1.52, 1.12, 0.64, and 0.95 for **c** to **f**, respectively.

### 9 Comparing the SNR of experimental measurements

We estimate the SNR value of the experimental measurements by computing

$$\begin{aligned} I_{max} &= \max\left(I - I_{background}\right), \\ \text{SNR} &= \frac{I_{max}}{\sqrt{I_{max}}} = \sqrt{I_{max}}, \end{aligned} \tag{11}$$

where $I$ is the intensity trace (of individual pixel of an input image stack), and $I_{background}$ is minimum intensity of the input image stack (i.e. the ROI), which is used as the background value. $I_{max}$ is the maximum value above the background value. $I_{max}$ is an estimate of the photons emitted by the particles. It is assumed to follow a Poisson distribution and the variance of the signal is therefore $I_{max}$. We use $I_{max}$ to estimate the maximum SNR achievable in each sample.

For a fair comparison, we evaluate an ROI of the first 2,500 frames 6×6 pixels for all the experimental measurements. For the drosophila measurement, we used the same regions in Supplementary Fig. 7.1a. For CHO-K1 PMT-mECFP measurement, we used the upper left 6×6 region of Supplementary Fig. 6.1b. Similarly, we used the upper left 6×6 region for the 2,500-frame HeLa GPI-GFP measurements at different laser power in Supplementary Fig. 8.1, one DOPC lipid bilayer measurement in Fig. 2a, and one in-focused 100 nm bead measurement in Fig. 2c. We compute the SNR value for each pixel of the image stack using equation (11) and we show the box plot of the values in Supplementary Fig. 9.1.

Supplementary Fig. 9.1 shows the SNR box plots in log scale. Comparing the median of the SNR values, the 100 nm bead solution is about 2.0× and 1.6× than that of the drosophila measurements between nuclei and within a nucleus, respectively. In the 2D case, the median SNR value of DOPC lipid bilayer Rhod PE at 11 $W/cm^2$ is 1.4×, 2.0×, 1.7×, 1.4×, and 1.4× compared to the median SNR values of CHO-K1 PMT-mEGFP 3.8 $W/cm^2$, HeLa GPI-GFP 0.4 $W/cm^2$, HeLa GPI-GFP 1.0 $W/cm^2$, HeLa GPI-GFP 1.8 $W/cm^2$, and HeLa GPI-GFP 4.0 $W/cm^2$, respectively. Furthermore, we observe an increase in SNR as we increase the laser power in the HeLa GPI-GFP measurements. The trend matches the improvement in *ImFCSNet* predictions as seen in Supplementary Figs. 8.1c–f.

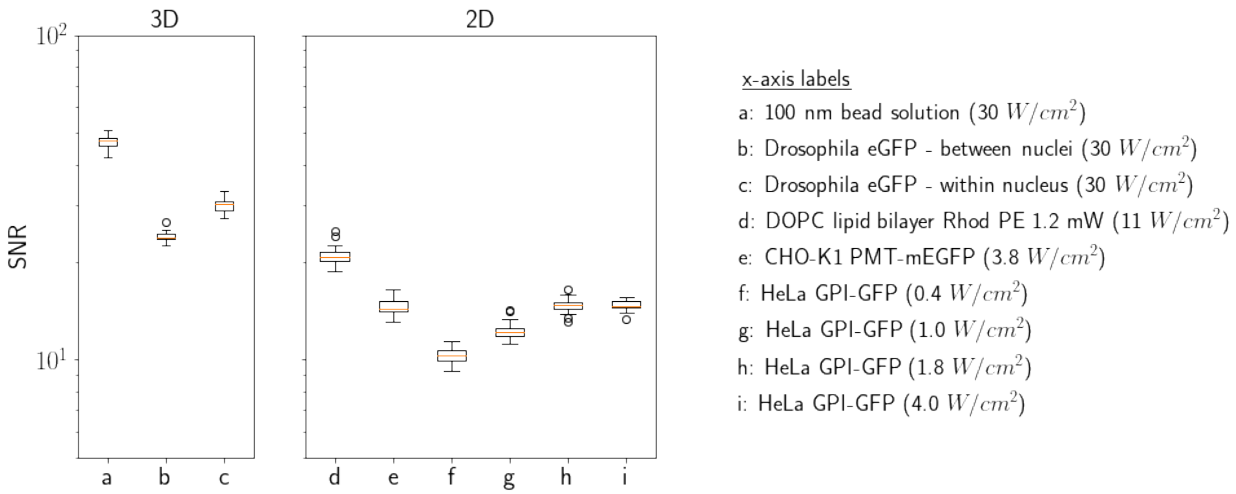


**Supplementary Figure 9.1 : Comparing the SNR of different experimental measurements.** In the 3D case, the laser power used was 30 $W/cm^2$. The median of SNR values of 100 nm bead measurement is 2.0× than that of drosophila eGFP – between nuclei measurement and 1.6× of drosophila eGFP – within a nucleus. In the 2D case, the median of SNR values of DOPC lipid bilayer Rhod PE 11 $W/cm^2$ is 1.4×, 2.0×, 1.7×, 1.4×, and 1.4× compared to the medians of CHO-K1 PMT-mEGFP 3.8 $W/cm^2$, HeLa GPI-GFP 0.4 $W/cm^2$, HeLa GPI-GFP 1.0 $W/cm^2$, HeLa GPI-GFP 1.8 $W/cm^2$, and HeLa GPI-GFP 4.0 $W/cm^2$, respectively.
